# A fully phased interspecific grapevine rootstock genome sequence representing *V. riparia* and *V. cinerea* and allele-aware annotation of the phylloxera resistance locus *Rdv1*

**DOI:** 10.1101/2022.07.07.499180

**Authors:** Bianca Frommer, Ludger Hausmann, Daniela Holtgräwe, Prisca Viehöver, Bruno Hüttel, Richard Reinhardt, Reinhard Töpfer, Bernd Weisshaar

**Affiliations:** Bielefeld University, Chair of Genetics and Genomics of Plants, Faculty of Biology & Center for Biotechnology (CeBiTec), Bielefeld, Germany; Julius Kuehn Institute (JKI), Institute for Grapevine Breeding Geilweilerhof, Siebeldingen, Germany; Max Planck Genome Centre Cologne, Max Planck Institute for Plant Breeding Research, Cologne, Germany

**Keywords:** Assembly, PacBio, SMRT sequencing, grapevine, rootstock, genome, Canu, annotation, resistance, resistance gene analogs, phylloxera, Rdv1

## Abstract

The phylloxera resistant rootstock cultivar ‘Börner’ is an interspecific hybrid derived from *Vitis riparia* and *V. cinerea* and a valuable resource for *Vitis* disease resistances. We created a fully phased, high-quality ‘Börner’ genome sequence named BoeRC using long PacBio reads. Comprehensive gene annotation of both ‘Börner’ haplotypes, designated BoeRip and BoeCin, was applied to describe the phylloxera resistance locus *Rdv1*. Using a mapping population derived from a susceptible *V. vinifera* breeding line and ‘Börner’, the *Rdv1* locus was further delimited. *Rdv1*, which is derived from *V. cinerea* and included in the haplotype BoeCin, was compared with sequences of phylloxera-susceptible and phylloxera-tolerant cultivars. Between flanking regions that display high synteny, we detected and precisely characterized a diverse sequence region that covers between 202 to 403 kbp in different haplotypes. In BoeCin, five putative disease resistance genes were identified that represent likely candidates for conferring resistance to phylloxera.

## Background

*Vitis vinifera* Linné subspecies *vinifera* is the most cultivated *Vitis* species worldwide. It is economically highly important and appreciated for wine production, table grapes, dry fruits like raisins, and other products. The species is endemic to Europe, and domestication and development of cultivars has a very long history ^1^. *V. vinifera* is highly susceptible to many diseases and pests including downy and powdery mildew as well as to the root aphid phylloxera (*Daktulosphaira vitifoliae* Fitch) ^2^. In the 19^th^ century, phylloxera caused a crisis of unprecedented dimension for viticulture and questioned the survival of viticulture in general. *Vitis vinifera* plants died quickly after phylloxera attack of the roots and huge cultivation areas were destroyed. Only the grafting of susceptible *V. vinifera* cultivars as scions to phylloxera-tolerant rootstocks derived from American wild *Vitis* species or their interspecific hybrids rescued viticulture ^3, 4^. The phylloxera plague remains as a serious case even if it is controlled at the moment.

The selection of a good rootstock that provides resistances or tolerances to the almost ubiquitous phylloxera is crucial for nowadays viticulture. One newly developed rootstock variety is the cultivar ‘Börner’, an interspecific F1 hybrid derived from a cross of the American *Vitis* genotypes *Vitis riparia* GM183 and *Vitis cinerea* Arnold. ‘Börner’ inherited full resistance to phylloxera on roots from *V. cinerea* and develops no nodosities or tuberosities ^5^ upon infestation with phylloxera (Fig. 1). *V. cinerea* is an American wild species known to often provide high resistance at the root and very high resistance at the leaf against phylloxera ^6^. A quantitative trait locus (QTL), located on chromosome (chr) 13, was identified to provide phylloxera resistance ^6^ and is referred to as *Rdv1* (resistance to *D. vitifoliae*)^7^. In addition, ‘Börner’ inherited positive viticultural traits including tolerance to drought, resistance to black rot caused by the fungus *Guignardia bidwellii* ^8^, and resistance to downy mildew caused by *Plasmopara viticola* ^9, 10^.

**Fig. 1:**
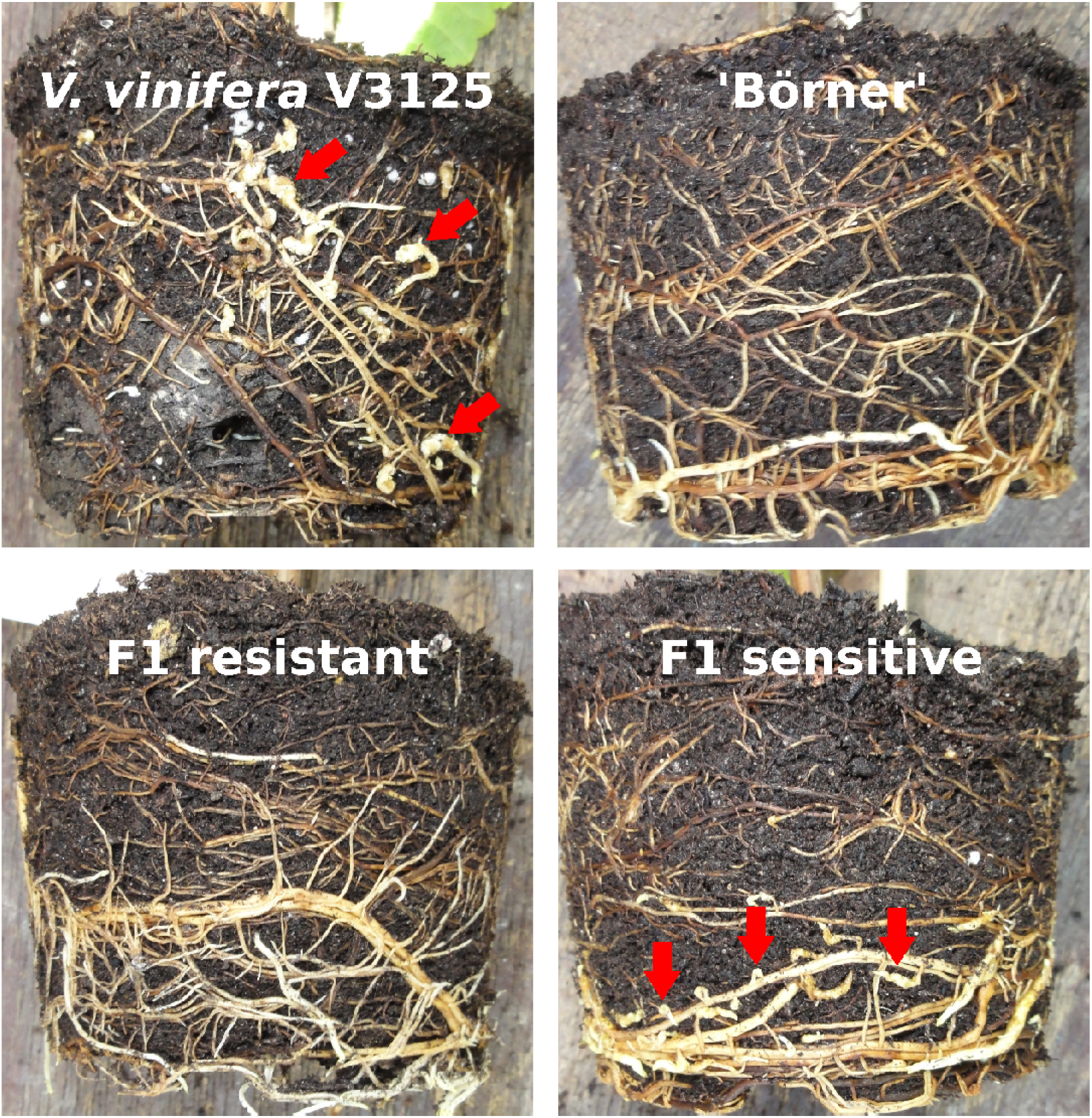
Root balls of susceptible and resistant grapevine plants attacked by phylloxera. Some phylloxera-induced nodosities on the roots are marked with red arrows. The pictures show roots of the grapevines indicated, the two F1 plants selected with the phenotypes as indicated are derived from the mapping population V3125 x ‘Börner’ ^6^.

Disease resistance, usually conferred through resistance genes, makes plants capable of surviving attacks from a broad variety of pathogens and pests ^11^. So called resistance gene analogues (RGAs) are identified by the protein domains they encode and structural features which are similar to validated resistance genes ^12^. Based on the protein domain structure that also indicates the localisation and/or site of action within the cell, they are broadly classified as nucleotide-binding site and leucine rich repeat domain receptor (NLRs) or pattern-recognition receptors (PRRs) ^11^. Usually, PRRs act at the cell surface because of a transmembrane (TM) domain, while NLRs cover a nucleotide-binding site (NBS) domain and usually act intracellularly.

*Vitis* genomes cover a haploid set of 19 chromosomes, and the haploid genome size varies around 500 Mbp ^13, 14^. The inbred cultivar ‘PN40024’ has been sequenced ^15, 16^ and the resulting assembly serves as a reference genome sequence (referred to as PN40024). However, since *Vitis* species and cultivars like almost all perennial plants are highly heterozygous and genetically divers ^10, 17, 18^, there is a need for more *Vitis* genome sequences. Several additional *Vitis* genome sequences have become available which were generated from long reads, including those of Cabernet Sauvignon ^19, 20^, Carménère^21^, Chardonnay ^14, 22^, *Muscadinia rotundifolia* cultivar Trayshed ^23^ and *V. riparia* Gloire de Montpellier (referred to here as VitRGM) ^24^. These assemblies each cover a primary and an alternate pseudo-haplotype of which the alternate pseudo-haplotype is usually of reduced contiguity. The cultivar *V. riparia* Gloire de Montpellier is tolerant against root infestation by phylloxera ^25^, but to the best of our knowledge it is unknown if this tolerance is linked to the *Rdv1* locus or to another region of the genome.

Long read DNA sequencing technology, like “Single Molecule, Real Time” (SMRT) sequencing ^26^ provided by Pacific Biosciences (PacBio), is one option to generate high quality genome sequence assemblies. The long reads more likely span problematic genomic regions and thus pave the way for more contiguous assemblies ^27^. Also, information of the haplotype and phase differences are more completely contained in a single read. This fact, together with the transmission of exactly one haplotype from two parents to a single offspring according to Mendel’s laws, is exploited by the “trio binning” approach for generating fully phased genome sequence assemblies ^28^.

To access and resolve the *Rdv1* locus, we set out to generate a haplotype-resolved genome sequence of ‘Börner’ with the goal to overcome complications caused by high heterozygosity and the complexity of resistance gene clusters. Using SMRT sequencing to generate long reads and additional Illumina short read data from both parents of ‘Börner’, we assembled both haplotype sequences of ‘Börner’ at chromosome level. The two truly phased haplotypes of the diploid interspecific hybrid ‘Börner’ represent the genome sequences of the two species *V. riparia* and *V. cinerea*. Structural and functional gene annotation was performed for both haplotype sequences, and RGAs were studied. The new sequence and annotation data as well as additional mapping results were used to dissect the *Rdv1* locus of ‘Börner’ at the gene level.

## Results

### Sequencing data of ‘Börner’ and its parents

To assemble the ‘Börner’ genome sequence, ~66 Gbp of raw SMRT sequencing data comprising 7,328,737 subreads with an average read length of 9,016 bp and an N50 length of 12,963 bp were generated. The expected diploid genome size (2n) of ‘Börner’ is 2×500 Mbp; thus the calculated coverage with long reads for a genome sequence with merged haplotypes would be 132 times, or more than 60-fold coverage for each individual haplotype. To allow k-mer-based binning of the long reads according to the two haplotypes that ‘Börner’ inherited from its parents, the parental genotypes were sequenced with Illumina technology to about 140-fold coverage.

### A ‘Börner’ genome sequence assembly in two phases

To generate a ‘Börner’ genome sequence with two separated haplotypes, the trio-binning approach was applied ^28^. The two haplotype-specific ‘Börner’ read subsets or bins, designated Vrip and Vcin, contain approximately 50 % of all reads and bases. These were assembled with Canu (see Methods). Statistics of the two resulting haplotype assemblies are shown in Table 1. Both assemblies reached the expected genome size of ~500 Mbp. The BoeRip and BoeCin haplotype assemblies represent “1n double haploid” genome sequences of the two species *V. riparia* and *V. cinerea*, respectively.

**Table 1.** Assembly statistics for the raw and scaffolded haplotype assemblies BoeRip and BoeCin of ‘Börner’.

|  | Scaffold Level<sup>a</sup> |  | Total |  |
| --- | --- | --- | --- | --- |
|  | BoeRip | BoeCin | Contigs<sup>b</sup> | Scaffolds |
| Sequences | 971 | 767 | 1,843 | 1,738 |
| Size (bp) | 495,882,484 | 501,563,116 | 997,019,452 | 997,445,600 |
| Largest seq. (bp) | 14,614,105 | 16,784,012 | 16,784,012 | 16,784,012 |
| N50 (Mbp) | 5.29 | 5.34 | 4.79<sup>c</sup> | 5.29 |
| N90 (Mbp) | 0.72 | 0.51 | 0.49<sup>c</sup> | 0.55 |

<sup>a</sup>scaffolded haplotype assemblies
<sup>b</sup>raw haplotype assemblies, numbers include contigs shorter than 10 kbp
<sup>c</sup>see Supplementary File 1 Fig. S1 for details

### Pseudochromosome construction and validation by genetic map

The assignment of scaffolds to pseudochromosomes was mainly based on the PN40024 reference sequence and was achieved by using RBHs (see Methods). For BoeRip, 446 scaffolds accounting for 95.21 % of all bases were assigned to pseudo-chromosomes. For BoeCin, 320 scaffolds holding 93.89 % of all bases were assigned. Since the reference contains a pseudochromosome chr00 for unassigned sequences, this has been created as well and is referred to as “chrUn”. In addition, some scaffolds find no clear corresponding sequence in the reference and result in a pseudochromosome “chrNh” (for no hit, see Fig. 2A). As a result, the haplotype assemblies each consist of 21 pseudochromosome sequences representing the 19 true chromosomes and two artificial sequences.

**Fig. 2.**
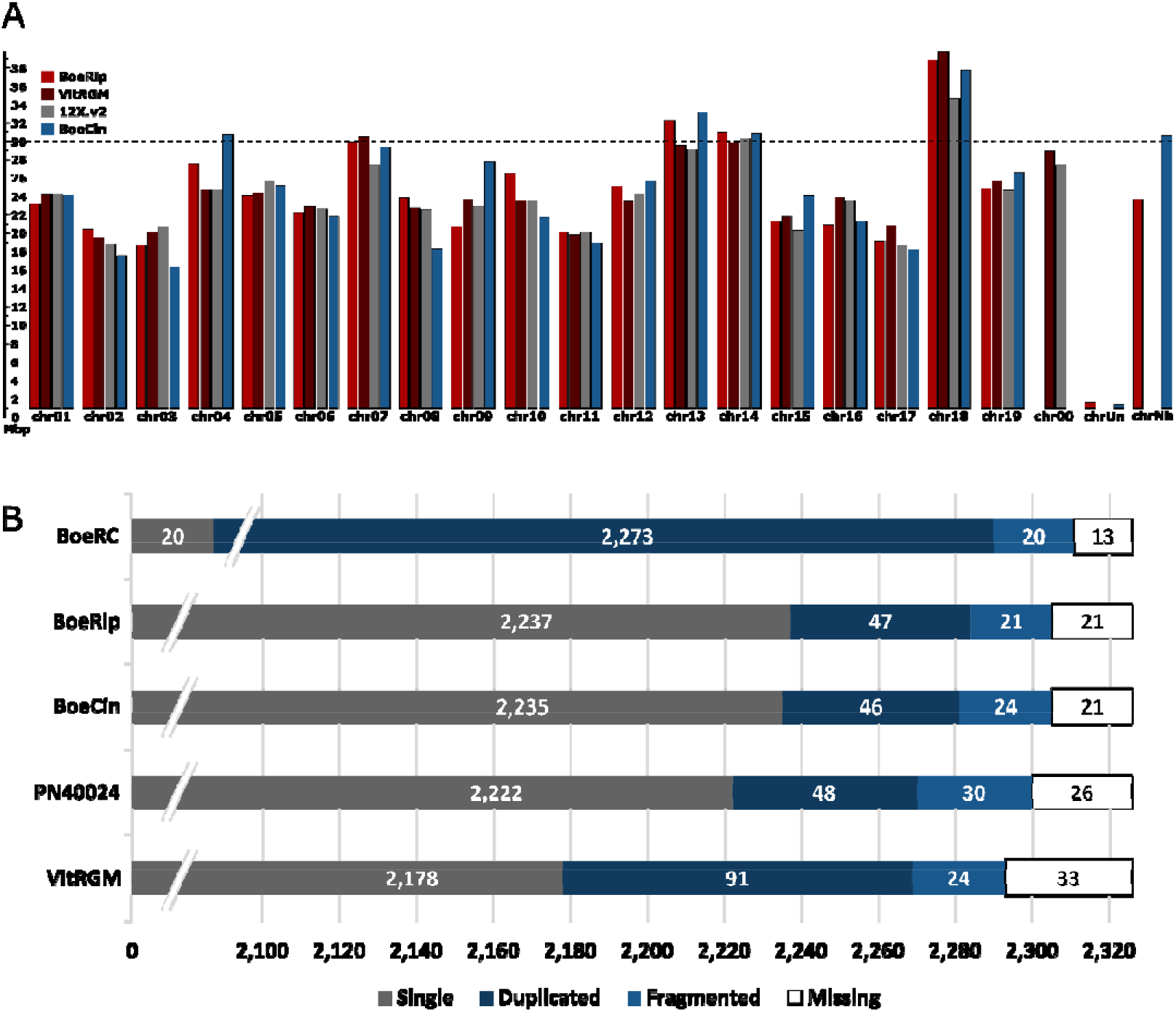
Comparison of the ‘Börner’ genome sequence assembly with other *Vitis* assemblies. **(A)** Pseudochromosome lengths comparison of both ‘Börner’ haplotypes, PN40024 and VitRGM. The bars represent the pseudochromosomes of the different assemblies. Red, BoeRip; dark red, VitRGM; grey, PN40024; blue, BoeCin. ChrUn holds sequences that were assigned to chr00 of PN40024 or VitRGM, and chrNh collects sequences with no assignment (Supplementary File 2 Table S1). **(B)** Plant core gene content (2,326 eudicots genes in reference set) of the assemblies in comparison to the PN40024 and VitRGM genome assemblies. Note that the bar graph is truncated at the left and focusses on only the duplicated, fragmented and missing BUSCO genes. The track labelled BoeRC represents the whole ‘Börner’ assembly (combined BoeRip and BoeCin sequences; merged results from the two haplotypes).

To validate the pseudochromosomes, 314 SSR markers that show unequivocal chromosome mappings on the haplotype assemblies were evaluated relative to their position on PN40024. A total of 309 markers confirm the pseudochromosome assignment. Conflicting markers were checked manually and corrected if possible (see Methods). The completeness of the assemblies was evaluated by detection of plant core genes with BUSCO (Fig. 2B). For both BoeRip and BoeCin, about 96.5 % of the conserved single copy gene set were detected. This indicated that both haplotype assemblies are more complete than the assemblies PN40024 and VitRGM (Supplementary File 2 Table S2).

### Evaluation of assembly quality and phasing

For validation of the phasing, bacterial artificial chromosome (BAC) sequences (Supplementary File 2 Table S3) with known haplotype/parental origin were mapped on BoeRip and BoeCin (Supplementary File 2 Table S4). The BACs were selected to cover regions on chr01 and chr14 and were sequenced with a read coverage of about 1,000-fold (Supplementary File 2 Table S5). BAC contigs of the same haplotype map with almost no sequence difference, while BAC contigs derived from the other haplotype display a wide range of mismatches and InDels (Supplementary File 1 Fig. S2). On both haplotypes, ~3.6 Mbp (chr01) and ~5.5 Mbp (chr14) were covered by BAC contigs of both phases and support a correct phasing over 9 Mbp.

In addition, the assembly quality evaluation tool Merqury (phasing assessment for genome sequence assemblies) was used to assess the quality of both haplotype assemblies. The overall base quality value (QV, consensus quality value, representing the log-scaled probability of error for consensus base calls) of 37.42 found for BoeRC indicates a very high level of correctness (see Supplementary File 2 Table S2 for comparisons to other assemblies). It should be noted that Merqury requires Illumina data for the complete trio (both parents and F1 as available for ‘Börner’ and its ancestors) to calculate the complete set of quality values.

Also the phasing accuracy was evaluated with Merqury (Supplementary File 1 Fig. S3). Based on haplotype-specific k-mers (referred to as hap-mers in Mercury), 2,759 phase blocks with an N50 of 2.25 Mbp were calculated for BoeRip and 1,852 blocks with an N50 of 2.42 Mbp for BoeCin. Almost all bases were covered by a block and the switch error rate was estimated to be less than 0.05 % (Supplementary File 2 Table S6; Supplementary File 1 Fig. S4). Overall, the quality of the ‘Börner’ genome sequence as well as its separation into haplotypes derived from *V. riparia* GM183 (BoeRip) and *V. cinerea* Arnold (BoeCin) is considered to be high.

### Sequence similarities and variations between BoeRip, BoeCin, PN40024 and VitRGM

All-versus-all alignments between the pseudochromosomes of the two ‘Börner’ haplotypes, and between each haplotype and PN40024 as well as VitRGM, were computed and visualised as dot plots (Fig. 3, Supplementary File 1 Fig. S5 to S8). All homologous pseudochromosomes show strong synteny, yet some rearrangements and smaller and larger gaps indicating insertions, deletions and/or missing sequence were detected.

**Fig. 3.**
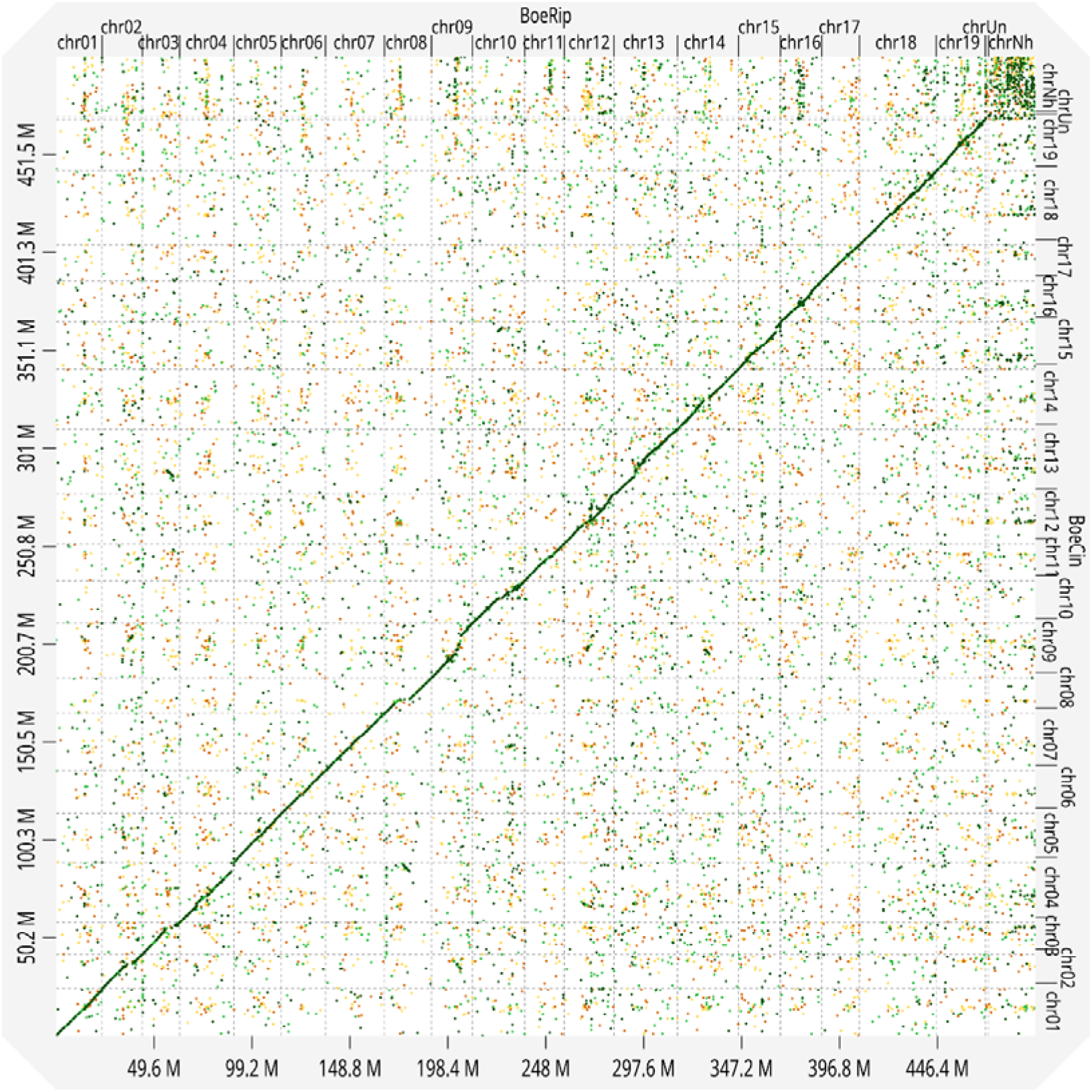
All-versus-all dot plot of both ‘Börner’ haplotypes. The graphic shows the dot plot between concatenated pseudochromosomes of BoeRip and BoeCin.

On average 90.20 % of all bases with an identity of 95.52 % were aligned between the pseudochromosomes of the two haplotypes except of chrUn and chrNh (see Methods). The SNP and InDel frequency for the protein coding fraction was 1/1,001 bp and 1/34 bp for the non-coding fraction of the genome sequence. When aligning with PN40024, on average 88.59 % and 87.51 % of all bases were aligned with an identity of 94.82 % (BoeRip) and 94.94 % (BoeCin) over all pseudochromosomes, respectively. The alignments resulted in a SNP and InDel frequency of 1/932 bp (BoeRip) and 1/935 bp (BoeCin) for the coding regions and of 1/32 bp for the non-coding regions.

An alignment of BoeRip and BoeCin with VitRGM resulted in 93.22 % (BoeRip) and 88.45 % (BoeCin) aligned bases with an identity of 98.53 % (BoeRip) and 95.29 % (BoeCin). The SNP and InDel frequency was 1/3,128 bp (BoeRip) and 1/1,019 bp (BoeCin) for coding regions and 1/94 bp (BoeRip) and 1/35 bp (BoeCin) for non-coding regions.

### Gene annotation

In total, ~274 Mbp (55.24 %, BoeRip) and ~278 Mbp (55.46 %, BoeCin) repetitive sequences were identified with the MAKER pipeline (Supplementary File 2 Table S7). The final gene annotation comprises 27,738 and 27,995 protein-coding genes (see Data availability for links to the genome sequence annotation of the *V. cinerea* and the *V. riparia* haplotypes) and 525 and 419 tRNA genes for BoeRip and BoeCin, respectively (Supplementary File 2 Table S7). Of the protein-coding genes, 19,500 genes of each haplotype were captured in RBHs. Based on RBHs, BoeRip shares 17,922 genes with VCost.v3 (PN40024) and 18,928 genes with VitRGM; BoeCin shares 17,837 genes with VCost.v3 (PN40024) and 18,126 genes with VitRGM.

### Resistance genes

The annotated genome sequence of ‘Börner’ was used to identify RGAs based on their typical protein domain structure. Resistance gene annotation with RGAugury revealed 2,081 RGAs for BoeRip and 2,009 RGAs for BoeCin (Fig. 4). Of these, 1,509 were classified as RBHs between BoeRip and BoeCin.

**Fig. 4.**
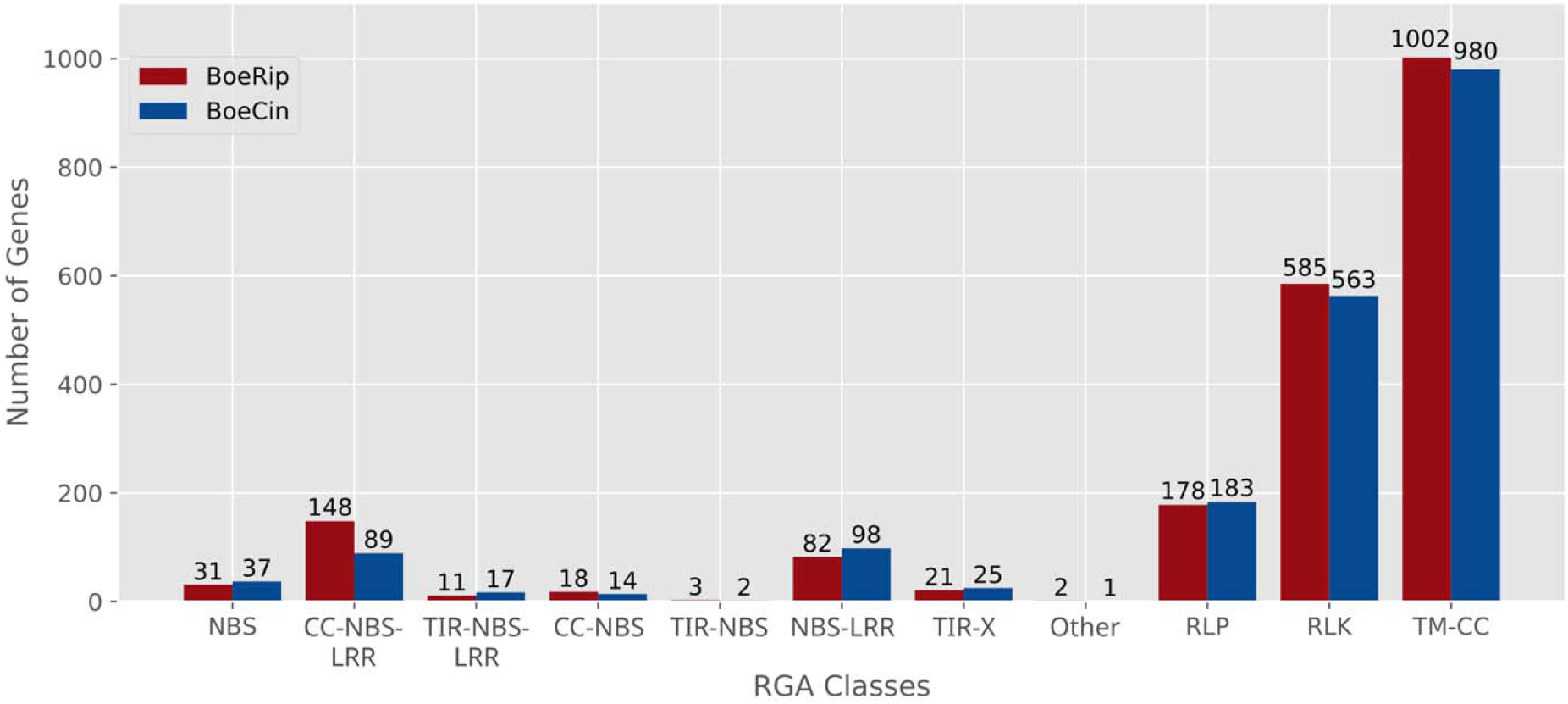
Distribution of genes to RGA classes based on characteristic protein domains. The red bars display the number of RGA genes of BoeRip and the blue bars the RGA genes of BoeCin. RGAs were classified according to the domains encoded by the predicted genes. Designations were adapted according to ^29^. NBS, Nucleotide Binding Site; CC-NBS-LRR with CC for Coiled-Coil and LRR for Leucine Rich Repeat; TIR-NBS-LRR with TIR for Toll/Interleukin-1 Receptor like; TIR-X with X for unknown domain; Other including TIR-CC-NBS-LRR or TIR-NBS-LRR-CC-TM with TM for TransMembrane region; RLP, Receptor Like Protein; RLK, Receptor Like Kinase.

### Candidate genes for Rdv1

At the sequence level, the region between GF13-01 and GF13-11 that contains *Rdv1* displays a highly diverged structure between BoeRip, BoeCin, PN40024 and VitRGM (Fig. 5A, Supplementary File 1 Fig. S9, S10).

**Fig. 5.**
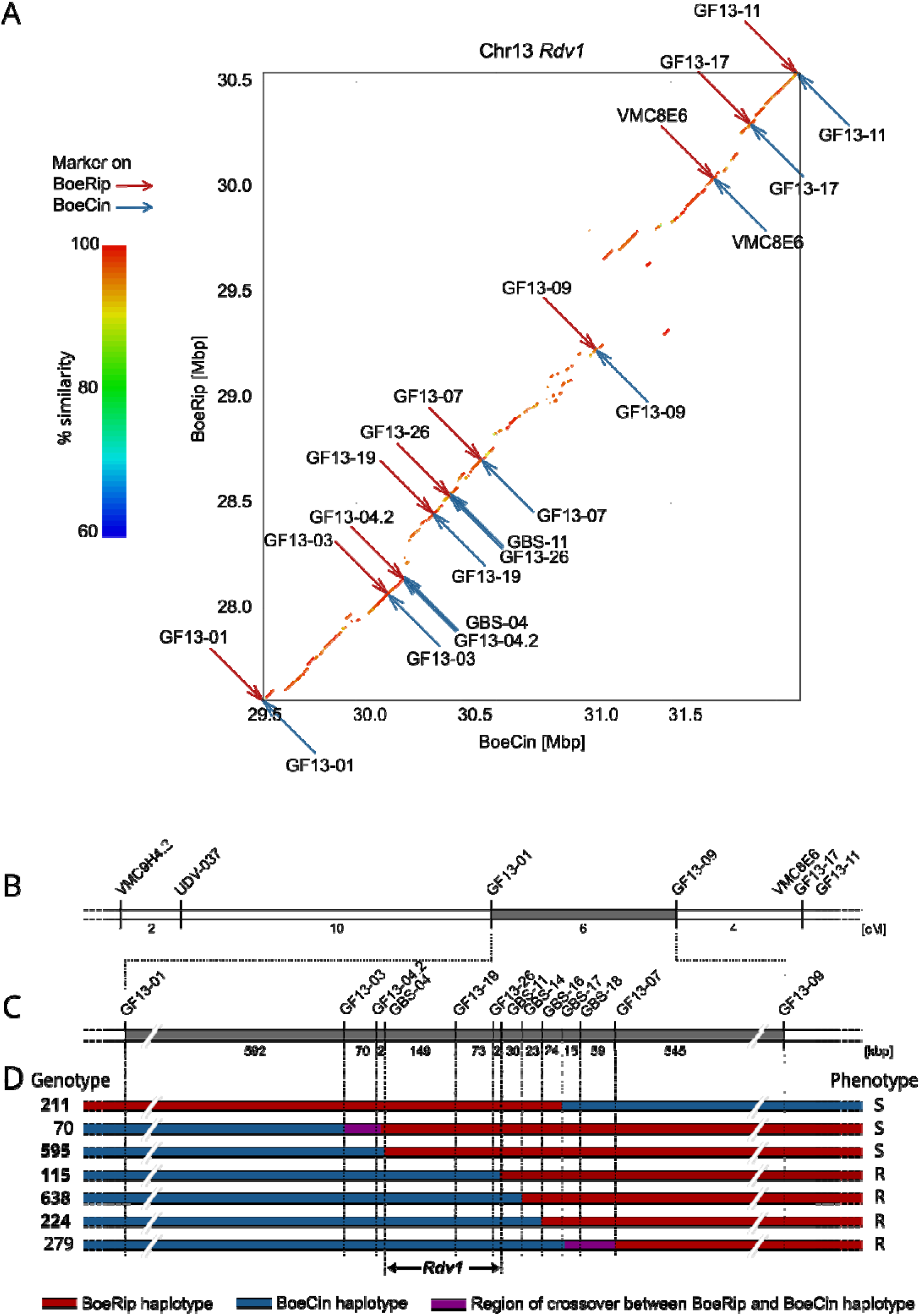
Integration of genetic map and haplotype-specific physical map of the *Rdv1* region on pseudochromosome 13. **(A)** Dot plot of the *Rdv1* region between BoeRip and BoeCin. Genetic markers mapping on BoeRip are represented as red arrows and markers mapping on BoeCin as blue arrows. The *Rdv1* locus is located between GBS-04 and GBS-11. (**B-D)** Fine mapping of the phylloxera resistance locus *Rdv1*. **(B)** Genomic region of the genetically mapped *Rdv1* locus on chromosome 13 of ‘Börner’. The grey bar represents the interval between the delimiting SSR markers GF13-01 and GF13-09 ^6^, the numbers indicate the genetic distance between adjacent markers in cM. **(C)** Physical position of the SSR and GBS markers used for fine mapping of this region. The bar depicts the enlarged region taken from (B). Numbers refer to the physical distance of adjacent markers in kbp based on BoeCin (haplotype conferring resistance). **(D)** Local map of seven relevant F1 individuals from the V3125 x ‘Börner’ population revealed after haplotype-specific genotyping, only the ‘Börner’ haplotypes are shown. F1 genotypes analysed by GBS are highlighted in bold. The phenotypes are indicated on the right side: R, resistant; S, susceptible. Red and blue bars indicate the BoeRip and BoeCin haplotypes, respectively. Regions with unlocalised recombination sites are shown in purple.

The interval flanked by these genetic markers includes 2,978,547 bp with 179 genes for BoeRip and 2,538,570 bp with 173 genes for BoeCin, according to the BoeRC assembly and its annotation, as well as 2,796,801 bp with 273 genes for PN40024 and 5,854,243 bp with 450 genes for VitRGM according to published data. The interval between GF13-01 and GF13-11 in VitRGM, which has been generated with a FAL-CON-Unzip and which is representing a pseudo-haplotype, is twice as large as in the other assemblies.

To further delimitate *Rdv1*, new genetic markers were designed and used to establish a local map. F1 individuals with different flanking haplotypes were selected from the mapping population V3125 x ‘Börner’, genotyped and tested for phylloxera resistance. Since marker assay design tuned out to be difficult for gene-level resolution, genotyping-by-sequencing (GBS) was applied to five crucial F1 genotypes. Mapping of SSR and GBS markers as well as the presence or absence of the resistance encoded by BoeCin led to delimitation of *Rdv1* to a region between the markers GBS-04 and GBS-11 (Fig. 5B-D, Supplementary File 2 Table S8). The GBS markers were used to precisely locate recombination sites that were identified between SSR markers. Since the exact positions of the GBS markers are unknown for PN40024 and VitRGM, the markers GF13-04.2 and GF13-26, located very close to the markers GBS-04 and GBS-11, were used to specify the interval size.

Based on the gene annotation data generated for the assemblies, the RGA studies and intensive manual curation (see Methods), the genetically delimited *Rdv1* locus was evaluated in detail at the gene level (Fig. 6). The locus has a size of 223,738 bp with 20 genes and eight potential resistance genes in BoeCin, a size of 402,698 bp with 25 genes and 13 resistance genes in BoeRip, a size of 202,484 bp with 15 genes and seven resistance genes in PN40024 and 208,610 bp with 16 genes and four resistance genes (all detected as RGAs) in VitRGM. Complete gene structures were counted as “normal” genes even if the gene belongs to a TE. The *Rdv1* locus is covered by a single contig in both ‘Börner’ haplotypes.

**Fig. 6.**
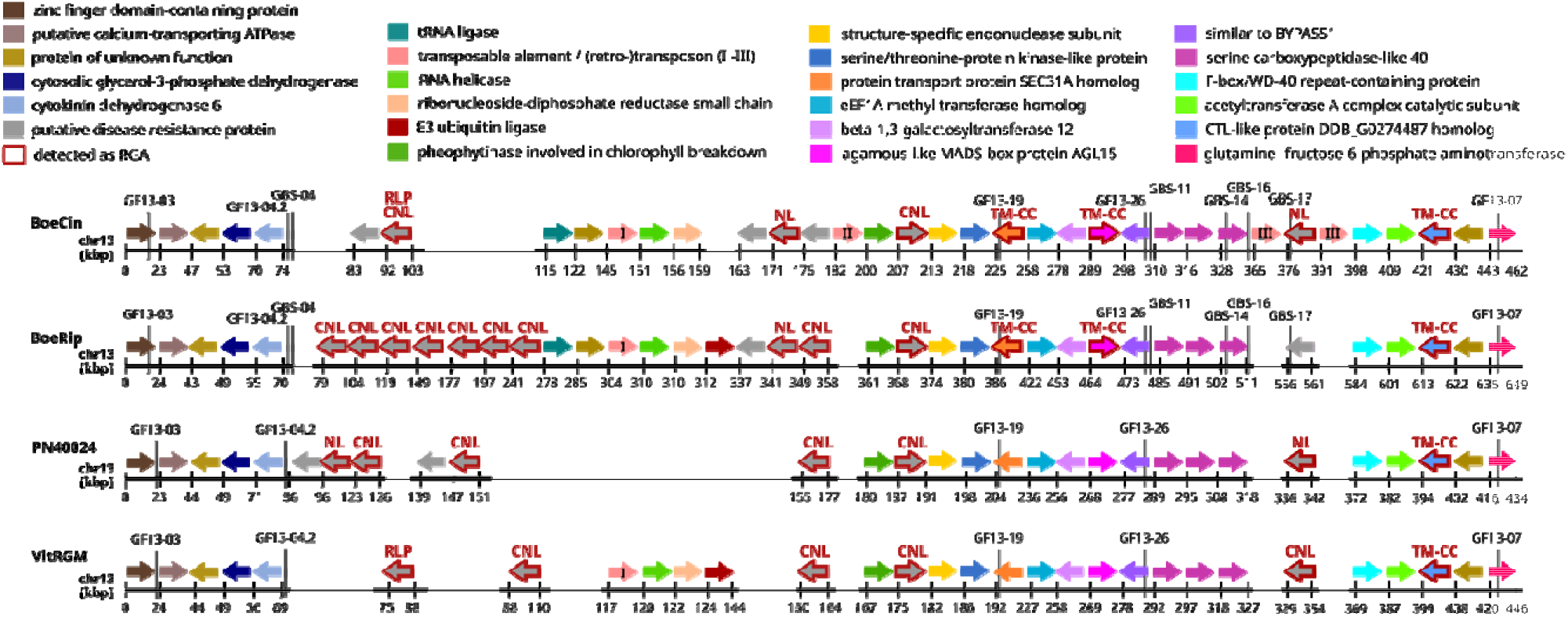
Genes of BoeCin, BoeRip, PN40024 and VitRGM at and surrounding the *Rdv1* locus. The genes (alleles) were clustered according to similarity and are shown in one colour for each orthogroup. The grey box represents the *Rdv1* locus delineated by the genetic markers GF13-04.2/GBS-04 and GF13-26/GBS-11; additional markers are included upstream and downstream of the *Rdv1* locus. The kbp values on the axis were adapted such that the start of the first gene is bp zero (relative coordinates). At the top, the colour code for each group of related genes/alleles and the corresponding functional annotation is given. Grey coloured genes are potential resistance genes according to their functional annotation; genes with a red edging were identified as RGA (see text). Above each RGA, the domain type classification of the encoded protein is mentioned. See legend to Fig. 4 for acronyms, CC-NBS-LRR abbreviated as CNL, NBS-LRR as NL. Three genes qualified as TE genes of different types, these are marked with roman numbers; I: mutator-like element (MULE), II: similar to transposon TX1 protein, III: similar to retrovirus-related polymerase polyprotein.

A resistance gene cluster that displays a very divergent structure and different gene content among the three analysed haplotypes and the pseudo-haplotype VitRGM is present between the cytokinin dehydrogenase encoding gene (dark blue in Fig. 6, annotation ID *BoeCin13g18380*, tagged by GF13-04.2/GBS-04) and the BYPASS1-related gene (purple in Fig. 6, annotation ID *BoeCin13g18411*, tagged by GF13-26/GBS-11). The haplotype conferring dominant resistance (BoeCin) contains eight potential resistance genes (Supplementary File 2 Table S9). Three of them, located in the southern part of the *Rdv1* locus (*BoeCin13g18393*, CC-NBS-LRR class; *BoeCin13g18396*, TM-CC class; and *BoeCin13g18399*, TM-CC class), are located in a syntenic region that contains similar alleles of the genes detected in all four haplotypes (see Discussion). In the following, the distinction between true haplotype sequences (PN40024, BoeRip and BoeCin) and the pseudo-haplotype VitRGM, that may contain merged sequences from the two phases of *V. riparia* Gloire de Montpellier, is neglected. Due to presence of these three genes in susceptible genotypes with highly similar alleles, they were considered not to be relevant for phylloxera resistance. The remaining five potential resistance genes (*BoeCin13g18381; BoeCin13g18382*, CC-NBS-LRR/RLP; *BoeCin13g18388; BoeCin13g18389*, NBS-LRR; *BoeCin13g18390*) can be considered as candidate genes for causing phylloxera resistance. *BoeCin13g18381, BoeCin13g18382* and *BoeCin13g18389* are putative disease resistance genes encoding RPP13-like proteins. The other two (*BoeCin13g18388*, *BoeCin13g18390*) are similar to the *Arabidopsis thaliana* resistance gene *AtLRRAC1*. To visualise the similarities between the protein-coding genes, especially the putative disease resistance genes among and within the haplotypes, a similarity matrix was calculated between BoeCin and itself as well the three other haplotypes (Supplementary File 2 Table S10 to S13). The similarities detected confirm the relations mentioned above, including the relatedness of, for example, the similarity of the putative disease resistance proteins encoded by *BoeCin13g18388* and *BoeCin13g18390*.

## Discussion

### Final ‘Börner’ genome sequence assembly

For the fully phased genome assembly, the sequence reads were binned into parental subsets prior to assembly. The binning process produced two main read sets, both containing approximately half of all reads. An additional small read set with relatively short reads that were probably derived from homozygous regions was also generated. The binning process provided an excellent basis for generating high quality, haplotype-separated genome assemblies, with sufficient sequence coverage. Two fully phased genome assemblies of the grapevine cultivar ‘Börner’ were computed. About 90 % of all bases were included in contigs equally or larger than 530 kbp (BoeRip) and 450 kbp (BoeCin). In contrast to other published grapevine genome assemblies, most of them generated with FALCON and FALCON-Unizp, the genome assembly size was not over- or underestimated by Canu and thus reaches for both haplotypes approximately 500 Mbp (Supplementary File 2 Table S2). As noted before ^14, 19, 24^, FALCON-Unzip sometimes overestimates the primary contigs and underestimates the haplotigs because it orders some alternative sequences of heterozygous regions to the primary contigs. Regarding genome assembly, TrioCanu/Canu ^28, 30^ turned out to be a very good choice for generating the genome sequence assembly BoeRC of the interspecific grapevine cultivar ‘Börner’. The N50 value of both ‘Börner’ haplotype assemblies outreaches most of the other *Vitis* genome assemblies. Comparing the N50, QV and overall k-mer completeness with some available chromosome level *Vitis* assemblies, BoeRC has with distance the highest N50 value, the highest quality values (estimated accuracy of 99.9998 %) and the highest degree of completeness (98.97 %) (Supplementary File 2 Table S2). To further demonstrate that the ‘Börner’ haplotype assemblies are fairly complete, the assemblies were investigated for the plant core genes. As in other high-quality genome sequence assemblies of grapevine like that of Cabernet Sauvignon ^20^, more than 98 % of complete core genes (BUSCOs) were identified.

### Phasing

The haplotype assemblies can be considered as fully and correctly phased. Through mapping of phased BAC sequences derived from parts of chr01 and chr14 of ‘Börner’, it was shown that several Mbp of chr01 and chr14 are truly phased. The haplotype-specific BAC sequences coincide with the correct haplotype and share plenty of mismatches with the other haplotype. Also, the mapping validated correctness of contiguity and chromosome assignment in those regions.

Additionally, the phasing was analysed through k-mer analysis with Merqury. The k-mer completeness of BoeRC over both haplotypes was rated to 99.04 % (BoeRip) and 99.51 % (BoeCin), while haplotype precision was estimated to 99.07 % (BoeRip) and 99.92 % (BoeCin) (Supplementary File 2 Table S2). Thus, the haplotype assemblies can be considered as completely and correctly phased. The purity of the haplotypes was also expressed through the very low switch error rate of 0.046 % (BoeRip) and 0.035 % (BoeCin) and the high block N50 value of 2.25 Mbp (BoeRip) and 2.42 Mbp (BoeCin). None of the scaffolds showed a notable amount of k-mers from the other haplotype (Supplementary File 1 Fig. S2).

There would potentially be room for improvement of the assemblies BoeRip and BoeCin by methods like chromosome conformation capture sequencing (Hi-C). Even if there are indications that the organization of chromosomes into territories within the nucleus of diploid organisms could allow to extract data supporting correct haplotype separation ^31^, we did not follow this route and anticipate that the quality of the BoeRC assembly is sufficient for almost any application with relevance to viticulture.

### Similarity of ‘Börner’ haplotypes and comparison with PN40024 and VitRGM

The assemblies BoeRip and BoeCin show overall similarity to each other and to PN40024. Even if the haplotypes among themselves have almost the same SNP frequency as with PN40024, more precisely 1 SNP in 33 bp compared to 1 SNP in 31 bp, more bases were aligned between the haplotypes with a higher identity than with PN40024. The difference in SNP frequency becomes more specific when comparing the SNP frequency of the coding regions. Here, BoeRip and BoeCin show less SNPs (1/1,001 bp) than with PN40024 (1/932 bp BoeRip & 1/935 bp BoeCin). Because of the high heterozygosity between haplotypes of grapevine ^32^, it is not surprising that the SNP frequency between ‘Börners’ haplotypes is almost the same as to another *Vitis* species. Reported SNP frequencies for grapevines range from 1 SNP per 64 bp ^33^ to 1 SNP per 200 bp ^17^. The SNP frequency between *V. riparia* and *V. vinifera* was determined to 1 SNP per 78 bp ^34^. Generally, it seems that SNP frequencies between and within *Vitis* species are not that different. However, the SNP frequency between BoeRip and VitRGM, both representing *V. riparia* genome sequences, was significantly lower than between BoeCin and VitRGM, for coding as well as non-coding regions. This may indicate that *V. riparia* GM183 and *V. riparia* Gloire de Montpellier were derived from related populations, or that the species *V. riparia* generally covers less variation.

High sequence identities and a good chromosome assignment were revealed through dot plots between homologous pseudochromosomes of the haplotypes of BoeRC and PN40024 or VitRGM. The “point clouds” often observed in the dot plots between homologous pseudochromosomes can be assumed to pinpoint the location of the centromere as centromeric regions are often highly repetitive and thus hard to sequence and assemble. The gaps detected in comparison with PN40024 may represent missing bases in the PN40024 genome sequence assembly. Almost all identified rearrangements between the haplotypes of BoeRC and PN40024 were found between both haplotypes and PN40024, and most of them are inversions. This calls for an improvement of the ‘PN40024’ reference genome sequence based on long read sequence data.

### Rdv1 locus

The *Rdv1* locus has been genetically mapped to a region of 224 kbp in size in the BoeCin haplotype of the ‘Börner’ genome. The genome regions orthologous to the *Rdv1* locus from BoeCin and different *Vitis* species are flanked on both sides by large syntenic regions. The high variability and divergence that is known for resistance gene clusters ^35^ was also detected within the *Rdv1* locus.

In syntenic regions, alleles may vary significantly in lengths between haplotypes, still they encode the same protein even among different *Vitis* species. For example, the PN40024 gene *Vitvi13g01613* which encodes a putative glycerol-3-phosphate dehydrogenase (dark blue gene in Fig. 6, northern part of the locus), is 3,600 bp longer than its BoeCin ortholog. The 466 amino acid long protein sequence is identical between all haplotypes analysed except one amino acid substitution each in PN40024 and BoeRip. Increased gene length of *Vitvi13g01613* in PN40024 is caused by a larger third intron. Several TE fragments and especially one larger LTR/Copia-like TE explain the difference in the length of intron three.

In addition to SNPs, quite some presence/absence variation (PAV) contributes to the divergence of the *Rdv1* region in *Vitis*. The various PAVs were, at least in several cases, explained by detection of TEs (either DNA transposons or retrotransposons) in the haplotypes (Fig. 6, Supplementary File 1 Fig. S11). For example, two intact LTR/Copia TEs and two intact DNA/hAT TEs were detected in the *Rdv1* region of BoeRip in the context of the tandem array of seven RGAs of the CC-NBS-LRR (CNL) type that is only present in BoeRip. Another genic PAV is the E3 ubiquitin ligase coding gene (*BoeRip13g18552*) that was detected in BoeRip and VitRGM which might be specific for *V. riparia*, although this hypothesis needs to be tested with more data from *V. riparia*. The resistance gene *BoeCin13g18388* encodes a LRR domain and is about 2,400 bp longer in BoeCin than in BoeRip (*BoeRip13g18553*). The reason is again an intron in the 5’UTR region of BoeCin which includes an LTR/Gypsy TE. This TE insertion causes a shifted translation start and the protein of the BoeCin allele is 49 amino acids longer at the N-terminus than the protein encoded by the BoeRip allele. In case of the MULE transposon (TE with similarity to mutator-like elements, marked with I in Fig. 6), a gene coding for a transposase protein was identified in BoeCin, BoeRip and VitRGM. There are hints from the TE annotation (Supplementary File 2 Table S7) that the MULE transposon might contain two ORFs in opposite orientation which is characteristic for some MULEs. However, the terminal inverted repeats were not identified. Because of the great diversity within the same and between transposon superfamilies, generally and also with regard to the structure and length of MULEs ^36, 37^, further work is needed to precisely identify all TEs and TE fragments that contribute to PAVs at this and other loci.

The *Rdv1* region is characterized by one extended resistance gene cluster. Orthologous regions in the other studied haplotypes derived from different *Vitis* species display very divers cluster structures with respect to length in kbp and number of resistance genes. In the BoeCin haplotype that confers the dominant resistance to phylloxera and which segregates in offspring of ‘Börner’, the cluster has a length of about 350 kbp and includes 10 potential resistance genes. The cluster extends at the southern end beyond the genetically delimited *Rdv1* locus. In the BoeRip haplotype that confers recessive susceptibility to phylloxera, the cluster includes 14 potential resistance genes.

In both ‘Börner’ haplotypes as well as in VitRGM, the cluster shows an insertion of a conserved array of four to six genes compared to PN40024 representing *V. vinifera* in the comparison. The inserted genes (or the genes deleted in PN40024) encode a tRNA ligase, a protein of unknown function, a mutator-like element (MULE) DNA transposon, a DEAD-box ATP-dependent RNA helicase, a ribonucleoside-diphosphate reductase small chain and a ubiquitin ligase. These genes display no features of resistance genes and are found with high identity values in the susceptible haplotype BoeRip. They are, therefore, no candidate genes for *Rdv1*.

Within the southern part of the locus, RGAugury detected two RGAs with TM-CC domains in the encoded proteins that are potential false positive results (orange and pink in Fig. 6, with red edging in BoeRip and BoeCin due to the RGAugury output). Based on the functional annotation (Supplementary File 2 Table S9), the two orthogroups code for homologs of SEC31A (a component of the coat protein complex II (COPII) involved in the formation of transport vesicles) and homologs of AGL15 (agamous-like MADS-box factor potentially involved in control of development). The genes in the syntenic block that starts at its northern end with the homolog of *SLX1* (encoding a subunit of a structure-specific endonuclease complex involved in processing diverse DNA damage intermediates, light ochre in Fig. 6) and which extends to the south beyond the delimiting markers GF13-04.2/GBS-04, are also very unlikely to be candidates for *Rdv1*.

To the north, the synteny extends for two more genes and terminates at a TE gene annotated as TX1-like non-LTR retrotransposon (*BoeCin13g18391*, marked II in Fig. 6) that is only detected in BoeCin. Southern to this TX1-like TE, a syntenic orthogroup (including *BoeCin13g18393* in BoeCin) encodes a CC-NBS-LRR (CNL) resistance protein related to the wild potato resistance protein RGA4. Since this gene is represented by closely related alleles in BoeCin and the haplotypes conferring susceptibility (amino acid sequence identity: BoeRip 99.9 %, PN40024 97.1 %), it is not considered to be a candidate for *Rdv1*.

Based on this evaluation, the most promising genes mediating phylloxera root resistance are the five resistance genes northern and southern to the array of four to six genes conserved in BoeCin, BoeRip and VitRGM (*BoeCin13g18381; BoeCin13g18382*, CNL/RLP; *BoeCin13g18388; BoeCin13g18389*, NL; *BoeCin13g18390*). Ideally, an F1 genotype from the V3125 x ‘Börner’ cross with a recombination site between the BoeCin and BoeRip haplotypes should have been identified. However, such an event was not detected. It is possible that the high divergence and sequence dissimilarity throughout the *Rdv1* locus causes a suppression or at least reduction of recombination events at the locus, which could explain this failure. A reduction of recombination frequency was, for example, also described for the *Rpv3.1* locus that confers resistance to *P. viticola* in *Vitis* ^38^.

The five remaining resistance genes can be divided into two types with respect to functional annotation. The resistance genes *BoeCin13g18388* and *BoeCin13g18390* were not detected as RGAs by RGAugury, but encode proteins similar to the *A. thaliana* resistance gene *AtLRRAC1* (*At3g14460*). *AtLRRAC1* has been reported to be involved in the defence against the biotrophic fungus *Golovinomyces orontii* as well as the hemi-biotrophic bacteria *Pseudomonas syringae*, and might be relevant for signalling via cAMP-dependent defence pathways ^39^. The three other resistance genes, namely *BoeCin13g18381, BoeCin13g18382* CC-NBS-LRR/RLP and *BoeCin13g18389* NBS-LRR (NL), encode proteins similar to *RPPL1* (Recognition of *Peronospora Parasitica* 13-Like 1). *AtRPPL1* (*At3g14470*) received its annotation from *AtRPP13* (*At3g46530*), a gene that confers resistance to the biotrophic oomycete *Peronospora parasitica* and which encodes an NBS-LRR protein ^40, 41^. In *A. thaliana, AtLRRAC1* and *AtRPP11* form a cluster of two neighbouring genes, a feature that is conserved in *Vitis* (Fig. 6). It will be interesting to figure out if the conserved co-localisation of two different resistance genes has functional implications for resistance to phylloxera in *Vitis* species.

In BoeCin and BoeRip, one of the resistance genes is classified as encoding an NBS-LRR (NL) protein. Despite an identical NBS domain with only very few substitutions, the LRR region varies greatly between the two alleles (*BoeCin13g18389* and *BoeRip13g18554*) and is shorter in BoeRip because of an early stop codon. Also for the other four candidate resistance genes, differences between the alleles in BoeCin, BoeRip and PN40024 were detected. For example, the protein encoded by *BoeCin13g18382* was detected as RGA based on CC-NBS-LRR domains in addition to features of an RLP. However, the genes detected at syntenic positions (alleles) of BoeRip and PN40024 lack these features either in part or fully.

Since the PN40024 sequence is derived from the most susceptible line in our comparison, the protein sequences of the five *Rdv1* candidate resistance genes of BoeCin were compared with all PN40024 protein sequences. No identical PN40024 protein sequence or proteins with identical resistance domain sequences were identified. Thus, the five BoeCin resistance genes are promising candidate genes for conferring resistance to phylloxera.

## Conclusions

The fully phase-separated genome sequence assembly BoeRC from the phylloxera-resistant rootstock ‘Börner’, and its structural and functional gene annotation, build a cornerstone for the investigation of loci from ‘Börner’ that are linked to various valuable traits. Here, the focus was on phylloxera resistance. The sequences of the haplotypes BoeCin conferring resistance to phylloxera and BoeRip conferring susceptibility allowed to precisely map recombination sites by GBS in crucial genotypes of a mapping population of V3125 x ‘Börner’. Subsequently, detailed examination and comparative genomics of the gene content of the *Rdv1* locus allowed to delimit the locus to a region of 123 kb in BoeCin haplotype with five resistance genes as potential candidates involved in mediating phylloxera resistance at the root. The resources generated will allow to study additional traits and have the potential to support resistance breeding for the benefit of viticulture.

## Methods

### Reference data sets

In this study, the ‘PN40024’ genome sequence ^16^ assembly 12X.v2 (urgi.versailles.inra.fr/Species/Vitis/Data-Sequences/Genome-sequences) and its annotation, the VCost.v3 gene annotation (https://urgi.versailles.inra.fr/Species/Vitis/Annotations) ^42^ as well as the *V. riparia* Gloire de Montpellier genome sequence assembly VitRGM and its annotation (https://ncbi.nlm.nih.gov/assembly/GCF_004353265.1) ^24^ were used for comparisons.

### Plant material and DNA and RNA extraction

‘Börner’ is listed in the *Vitis* International Variety Catalogue with the variety number *V*IVC 1499, its parents *V. riparia* GM183 and *V. cinerea* Arnold with variety numbers *V*IVC 4686 and *V*IVC 13645, respectively. All three cultivars do not belong to an endangered species and were obtained and are grown at the Institute for Grapevine Breeding Geilweilerhof at Siebeldingen (JKI Siebeldingen) in accordance with German legislation.

Dormant wood cuttings of the interspecific rootstock variety ‘Börner’ were raised in a growth chamber on soil at 18°-24°C (64°F-75°F), 70 % relative humidity and 16 h light and 8 h darkness. Prior to harvest the plants were cultivated in the dark for 72 hours. Small, young leaves were harvested and stored on ice until use. Extraction of high molecular DNA for SMRT sequencing was performed starting with 1.5 g fresh weight and was carried out with the NucleoSpin Plant II Maxi kit for DNA (Macherey and Nagel, Düren; Germany) according to the recommendations of the manufacturer for plants. DNA integrity, quality and quantity was checked using a TapeStation (Agilent) and was found to peak at a fragment size higher than 48.5 kbp.

Leaf material of ‘Börners’ parents was harvested from cuttings grown in the green house, frozen in liquid nitrogen and stored at −80°C. The DNA extraction was carried out with the DNeasy Plant Maxi Kit (Qiagen, Hilden; Germany) according to the instructions of the manufacturer. Tissue samples of leaves and tendrils of ‘Börner’ were collected from grown in the field. Total RNA was isolated using the Spectrum Plant RNA-Kit (Sigma, Taufkirchen; Germany).

Plant material for the local map of *Rdv1* was selected from the previously described mapping population V3125 x ‘Börner’ ^6, 43^ and another set of 310 F1 genotypes obtained after repeating the cross in 2006. To search for recombinant F1 lines, SSR markers associated with *Rdv1* (Supplementary File 2 Table S14) were used for genotyping as described ^43^. For GBS, young leaf material was harvested from selected individual F1 genotypes as well as from the maternal genotype V3125 and used for DNA extraction. Genomic DNA was prepared using an established CTAB-based protocol ^44^. Subsequently, the DNA obtained was treated with RNAse and quantified using PicoGreen.

### Library construction for ‘Börner’ genome and transcriptome sequencing

PacBio Sequel libraries were prepared according to the SMRTbell Template Prep Kit 1.0 and then size-selected with BluePippin for a target insert size of greater than 10 kb. Seventeen 1Mv3 SMRT cells were run on a Sequel I sequencer using the Binding Kit 1.0 and the Sequencing Chemistry version 1.0 (all from PacBio) for six hours per cell. The output BAM files of the sequencing step were loaded into SMRT Link 5.0.1 according to the recommendation of the manufacturer (PacBio Reference Guide 2018) and converted to FASTA format for downstream processing. The length distribution over all reads was calculated with the assembler Canu ^30^ and is provided in Supplementary File 1 Fig. S12.

The RNA samples from leaves and tendrils of ‘Börner’ were paired-end sequenced using Illumina technology on a HiSeq-2000 essentially as described ^45^. The new RNA-Seq read data (ENA/SRA study accession no. PRJEB46079) were processed together with data submitted before ^45^. These already existing RNA-Seq reads of ‘Börner’ (PRJEB34983) cover leaves (ERR3894001), senescent leaves (winter leaves, ERR3895010), inflorescences (ERR3894002), tendrils (ERR3894003) and roots (ERR3895007). Read data were trimmed with Trimmomatic-0.38 ^46^ allowing reads larger than 80 nt (new data) and 90 nt (published data) to be kept. The trimming statistics of the RNA-Seq data is provided in Supplementary File 2 Table S15.

### Illumina sequencing of the parental genomes of ‘Börner’

Library preparation for the *V. riparia* GM183 and *V. cinerea* Arnold samples was performed according to the Illumina TruSeq DNA Sample Preparation v2 Guide. Genomic DNA (1500 ng each) was fragmented by nebulisation. After end repair and A-tailing, individual adaptors were ligated to the fragments for PE sequencing. The adaptor ligated fragments were purified on a 2 % low melt agarose gel and size selected by excising a band ranging from 500-800 bp. After enrichment PCR of DNA fragments that carry adaptors on both ends, the final libraries were quantified by using the Quant-iT PicoGreen dsDNA assay on a FLUOstar Optima Platereader (BMG Labtech) and qualified on a BioAnalyzer High Sensitivity DNA Chip (Agilent). The *V. riparia* GM183 library was sequenced on a MiSeq PE run (2 × 250 nt) and on two lanes of a 2 × 100 PE run on a HiSeq-1500 in high output mode. The *V. cinerea* Arnold library was sequenced in one 2 × 150 bp PE run on a HiSeq-1500 in rapid mode. The read data were submitted to ENA (ENA/SRA study accession no. PRJEB45595). All genomic short read data were quality trimmed with Trimmomatic-0.36 ^46^ allowing reads equal or longer 80 nt to be kept. The trimming statistics is provided in Supplementary File 2 Table S16.

### Sequencing of pools of BACs from a BAC library of ‘Börner’

For the creation of phased sequences of ‘Börner’, 8 pools of BAC constructs containing a total of 440 mapped BACs from two loci (one on chr01, the other on chr14) were selected from a ‘Börner’ BAC library ^10^. The BACs were selected and distributed to the pools according to the positions of BAC end sequences on the corresponding regions of PN40024. Mapping to the reference assembly was carried out with the CLC Genomics Workbench v8.0 toolkit (https://www.qiagenbioinformatics.com/). Pools were arranged by excluding overlap of BAC inserts to allow assembly of individual BAC insert sequences. The BAC Pools1-4 were sequenced on a 454 Life Sciences Genome Sequencer GS FLX and PE (2 × 250 nt) on the Illumina MiSeq platform. The BAC Pools5-8 were sequenced on the MiSeq platform only. The read data were submitted to ENA (ENA/SRA study accession no. PRJEB46081). The MiSeq raw data were quality trimmed with Trimmomatic-0.32 using ‘ILLUMINACLIP: 2:40:15 LEADING:3 TRAILING:3 SLIDINGWINDOW:4:15 MINLEN:36’ ^46^. The trimming statistics for the BAC pools is provided in Supplementary File 2 Table S5.

### Library construction and sequencing for GBS

The barcoded libraries were prepared using Illumina library preparation kits using TrueSeq technology and sequenced on a NextSeq500. Sequencing was performed in 150 nt single-end (SE) modus and the read data were submitted to ENA. The ENA/SRA study accession no. is PRJEB53997. Information on genotypes, sequenc-ing strategy, data volume and run IDs are summarized in Supplementary File 2 Table S17.

### Phenotyping of phylloxera resistance

The test system used to assess phylloxera resistance of selected F1 individuals was essentially described previously ^6, 47^. The phylloxera population used for the artificial inoculation of potted plants was derived from leaves of naturally infested rootstock cultivars from the germplasm repository at JKI Siebeldingen. The phylloxera resistance tests started at the beginning of the vegetation season when enough infested leaf material was available as inoculum. Leaf pieces with 8 to 12 (3-5 mm) galls containing mature eggs were inserted in the soil of rooted cuttings of the F1 individuals. The inoculation procedure was repeated after 3 to 4 weeks. Approximately 4 weeks after the second inoculation the plants were rated as susceptible if nodosities were visible, or resistant if the roots developed normally. Each F1 genotype was tested in triplicate and the assays were repeated at least twice.

### Diploid genome sequence assembly

To compute a phased genome assembly, a binning and assembly approach was used. The binning prior to assembly was performed with the tool TrioCanu of the assembler Canu v1.7 ^28, 30^. TrioCanu uses short reads of the parents and long reads of the F1 as input. It bins the long reads of the F1 into parental subsets based on k-mer comparisons. The binning step was performed on a compute cluster using TrioCanu default parameters (k-mer size of 20). Binning resulted in one read subset for each parent and an additional small subset with reads that could not be assigned to either of the two parental haplotypes, referred to as UAR subset. In the following, the parental subsets are referred to as Vrip and Vcin read subset according to ‘Börners’ parents and the two different haplotypes combined into ‘Börner’, respectively. The UAR subset contained only 0.13 % of all bases with an average read length of 1.5 kbp.

After binning, the read subsets were used to compute the two haplotype assemblies BoeRip and BoeCin of ‘Börner’ with the assembler Canu v1.7 ^30^. To compute the BoeRip haplotype assembly, the Vrip and the UAR subsets were used in order to include identical and/or homozygous regions. Likewise, the BoeCin haplotype assembly was generated from the Vcin read subset and the UAR subset. Therefore, the basis of the haplotype assemblies, BoeRip and BoeCin, are approximately 32 to 34 Gbp read data with an estimated genome coverage of more than 60-fold each. For both separately computed haplotype assemblies, Canu was run on a compute cluster utilizing the parameters ‘genomeSize=550m’, ‘corMhapSensitivity=normal’, ‘correctedError-Rate=0.065, ‘canuIteration=1’ and ‘stopOnReadQuality=false’. During the assembly processes about 2.5 % of the corresponding input data (bp) remained unassembled. A search with blastn of the Basic Local Alignment Search Tool (BLAST) package+ v2.8.1 ^48, 49^ and the grapevine chloroplast ^50^ and mitochondrial sequences ^51^ was carried out on the haplotype assemblies and detected contigs were removed. Finally, the haplotype assemblies were polished twice with *arrow* v2.2.2 (smrtlink-release_5.1.0.26412) and the corresponding read subsets. Assembly statistics were computed with QUAST v4.6.3 ^52^.

### Scaffolding with ‘Börner’ BAC end sequences

To validate the quality of the two genome sequence assemblies, and also to scaffold a few smaller contigs, high-quality paired BAC end sequences from the ‘Börner’ genome (ENA/GenBank accession numbers KG622866 - KG692309 ^10^) were used. The 69,444 BAC end sequences cover 39,360,203 bp, have an average sequence length of 566 bp, and represent an eight-fold coverage of the ‘Börner’ genome based on average BAC insert length. First, the 62,498 BAC end sequences that represented pairs from one BAC were mapped to BoeRip and BoeCin with HISAT2 v2.1.0 ^53^ using the parameters ‘--maxins 2000000000’, ‘--secondary’, ‘--no-softclip’ and ‘--no-spliced-alignment’. Alignments with a mapping quality >10 were filtered out and alignment statistics were collected with SAMtools v1.8 ^54^. After mapping and quality filtering, the BAC end sequences were used to scaffold the haplotype assemblies with SSPACE (standard) v3.0 ^55^. SSPACE was run with the parameters ‘-x 0’, ‘-m 50’, ‘-k 3’, ‘-n 500’ and ‘-T 10’ and by using BWA-SW ^56^ for mapping. Contig extension was disabled (‘-x 0’) as the BAC end sequences are not phased and thus would eventually contaminate the haplotypes with a sequence from the other haplotype. However, a test run with enabled extension showed that the BAC end sequences were anyway not suitable for gap filling.

### Scaffold to pseudochromosome assignment

To build pseudochromosomes (the nucleotide sequence representation of chromosomes), an *ad hoc* gene prediction was generated with AUGUSTUS v3.3 ^57^. Prior to gene prediction, the RNA-Seq reads from ‘Börner’ (see above) were processed to create a *de novo* transcriptome assembly with Trinity v2.8.5 ^58^ with default settings. Hint files with information about exon and intron positions for the haplotype assemblies were created through mapping all trimmed RNA-Seq reads and the transcriptome assembly on the haplotype assemblies. The RNA-Seq reads were mapped with HISAT2 v2.1.0 ^59^ and soft-clipping disabled (‘--no-softclip’). The transcriptome sequences were mapped with BLAT v36×2 ^60^ and the parameters ‘-stepSize=5’ and ‘-minIdentity=93’. Hints were generated according to the AUGUSTUS documentation. Additionally, AUGUSTUS was trained with the CRIBI v2.1 gene annotation of PN40024 ^17^ to retrieve parameter sets for grapevine. Finally, genes were predicted on both haplotype assemblies BoeRip and BoeCin with the generated *Vitis* parameter set and the hint files.

The protein sequence of the primary transcript variant was extracted from the PN40024 CRIBI v2.1 annotation and from the *ad hoc* annotation of BoeRip and BoeCin. Reciprocal best BLAST hits (RBHs) between the protein sets of each of both haplotypes and the filtered proteins of PN40024 were computed with blastp utilizing an e-value threshold of 0.0001. For RBHs, two rounds of BLAST searches were performed. Here, one round was run with the proteins of one haplotype as input query and one with the PN40024 proteins as input query. Only RBHs with a percentage of identical matches and a query coverage per subject ≥80 % for at least one direction (e.g. PN40024 protein sequences as input query) were kept. Based on RBHs encoded, the scaffolds were assigned to the pseudochromosomes. Scaffolds with less than 10 RBHs in total and scaffolds that contain a significant number of RBHs linking a different pseudochromosome (30 % distance to the 2^nd^ rank in terms of RBH numbers required) were filtered out for later assignment or moved to the chromosomally unassigned part of the respective assembly.

Additionally, reciprocal hits between the scaffold sequences and the pseudochromo-some sequences of PN40024 were computed with blastn. The e-value, identity and query coverage filters remained the same as above. The nucleotide RBHs were iteratively computed and scaffolds assigned to pseudochromosomal positions. If the classifications based on protein and nucleotide level contradicted, the protein RBH classification was preferred.

To further refine the pseudochromosomes of the haplotypes, an additional assignment based on VitRGM was performed. Also, reciprocal hits between the scaffold sequences of one haplotype and the pseudochromosomes of the other haplotype and *vice versa* were calculated as described above. After manual construction of pseudochromosomes for the BoeRip and BoeCin phases, the refinement with the corresponding other haplotype was repeated until no further scaffolds could be assigned. The pseudochromosomes were constructed with gaps of 100 bp length between the scaffolds and with 10 kbp as minimal scaffold length. Due to the length filter, 61 (BoeRip) and 22 (BoeCin) scaffolds were discarded.

The orientation and order of the scaffolds on the pseudochromosomes was verified with dot plots and manually adapted if necessary. Thus, DNAdiff v1.3 of the MUM-mer package v4.0.0beta2 ^61^ was run pairwise on homologous pseudochromosomes of BoeRip or BoeCin and PN40024 with default settings and the resulting 1-to-1 alignments were visualized with mummerplot v3.5.

The pseudochromosomes were validated with 340 simple sequence repeat (SSR) genetic markers from ^43^ and the 4 SSR markers ATP3 ^62^, Gf13_11a, VMC8E6 ^6^ and Gf14-42 ^8^ (Supplementary File 2 Table S14). Primer sequences were mapped to the pseudochromosomes with primersearch of the EMBOSS v6.6.0.0 package ^63^ and with a blastn search. A total of 314 markers were assigned to unequivocal sequence positions, the remaining markers do not map, show non-evaluable multi-mappings or map with too many mismatches (Supplementary File 2 Table S18). Several markers did map only to one of the haplotypes (16 to only BoeRip and 30 to only BoeCin, see Supplementary File 2 Table S19). Emerging disagreements between the marker position on PN40024 and the BoeRip and BoeCin phases were further investigated through all-versus-all dot plots computed with the webtool D-Genies v1.2.0 ^64^ and by using long read mapping to detect sequence positions that are not or only very weakly supported by continuously mapping reads. The haplotype specific read subsets together with the unassigned read subset were mapped with minimap2 ^65^ v2.17 (‘-ax map-pb --secondary=no’) to the corresponding haplotype assembly and the coverage values per base were calculated with SAMtools. The haplotype assemblies are both covered at about 54x (Supplementary File 1 Fig. S13). The back-mapping results of the read subsets were consulted to reveal miss-assemblies if indicated by a genetic marker. A sequence location was further investigated if five reads end in a 10 bp region and if the read coverage five bp around the region drops to ≤ five.

Conflicting marker VVMD28 maps on chr03 of PN40024 and BoeCin, yet on chr13 of BoeRip. However, eight markers allocate the sequence to chr13 of BoeRip and neither a significant breakpoint was found in the read mappings nor in the all-versus-all dot plot (Supplementary File 1 Fig. S3). VCHR16B maps on chr16 of PN40024 and BoeCin, but on chr03 of BoeRip. Here, one SSR marker assigns the sequence to chr03 and no significant breakpoint was found in the read mapping or in the all-versus-all dot plot. The three SSR marker VVS4, VMC5G6.1 and UDV-126 map on chr08 of PN40024 and BoeRip, but on chr04 of BoeCin. The corresponding sequence was assigned to chr04 according to 376 protein RBHs, because only 141 protein RBHs assign it to chr08. Moreover, three SSR markers allocate the sequence to chr04, too. Through investigating the all-versus-all dot plot, an approximately four Mbp large alignment between chr04 and chr08 was found. However, since no position for a split was detected in the read mappings, the sequence remained on chr04 (Supplementary File 1 Fig. S6). GF10-06, VMC3D7 and UDV-073 mapped wrongly on chr02 of BoeRip instead of chr10; in this case the investigation of the respective contig resulted in a split of the sequence and a corrected assignment of the contig fragment in question. Due to 348 protein RBHs the sequence was initially assigned to chr02. The sequence showed the second most protein RBHs (171) with chr10. On marker side, 13 markers assign the sequence to chr02 and only three to chr10. However, the all-versus-all dot plot showed a large alignment between chr02 of BoeRip and chr10 of PN40024 (Supplementary File 1 Fig. S14), and the read mappings indicated a breakpoint position with a coverage drop to ≤ 5. Thus, the 12.3 Mbp sequence was split into a 7.7 Mbp large sequence that was assigned to chr10 and a 4.6 Mbp large sequence that remained at chr02. Another case was detected in the all-versus-all dot plots between chr13 of BoeCin and chr03 of PN40024 (Supplementary File 1 Fig. S6). The corresponding sequence was assigned to chr13 through 485 protein RBHs. Only 36 protein RBHs would allocate this sequence to chr03. Furthermore, 10 SSR marker support the assignment to chr13 and none to chr03. No coverage drop was found in the read mappings, the sequence remained on chr13 of BoeCin. Finally, only five genetic markers remained unresolved (VVMD28, VCHR16B, VVS4, VMC5G6.1, UDV-126).

To estimate the completeness of the assemblies, the plant core genes were localized with the program Benchmarking Universal Single-Copy Orthologs (BUSCO) v5.1.2 utilizing the database ‘eudicots_odb10’ (2,326 genes) ^66, 67^. For comparison, the BUSCOs of PN40024 and of VitRGM were also determined with identical parameters. The pseudochromosome lengths were visualized with cvit v1.2.1 ^68^.

### Validation of phasing

To validate the phasing of the ‘Börner’ haplotypes, the BAC sequence data of pool1-4 were *de novo* assembled with Newbler v2.6. The data of pool5-8 were assembled with CLC Genomics Workbench 8.0 using a word size of 30. Remaining vector sequences were removed with vectorstrip of the EMBOSS package v6.2. Assembled sequences shorter than 500 bp were discarded. The short reads of the parents *V. riparia* GM183 and *V. cinerea* Arnold were mapped with CLC’s map reads to reference algorithm to the assembled BAC sequences (linear gap cost, length & similarity fraction 1.0). Based on these mapping results regarding read coverage and percent of covered bases, the BAC sequences were assigned to either *V. riparia* GM183 or *V. cinerea* Arnold. BAC sequences with no read coverage were discarded. The assembled and phase-separated BAC contig sequences are available at https://doi.org/10.4119/unibi/2962639.

The ‘Börner’ BAC sequences with known haplotype allocation were mapped against the BoeRip and BoeCin haplotype assemblies with minimap2 v2.17 ^65^, sorted with SAMtools and the mappings visualized with the Integrative Genomics Viewer (IGV) v2.5.2 ^69, 70^. The number of base calls of the aligned BAC sequences on each haplotype were determined with SAMtools’ mpileup. Continuously and correct aligned BAC sequences show high base calls and low numbers of SNP/InDels (small nucleotide polymorphism, insertion-deletion polymorphism) to one of the two haplotypes and the opposite result with the alternative haplotype.

### k-mer based reference-free assembly evaluation with Mercury

Merqury v1.3 ^71^ was used to evaluate the diploid BoeRC assembly. First, the best k-mer size was determined with Merqurys best_k.sh script for the expected 500 Mbp haploid genome size resulting in 19-mers. Consequently, 19-mer databases were computed for the *V. riparia* GM183 and *V. cinerea* Arnold Illumina reads as well as for existing Illumina reads from ‘Börner’ ^10^. With these, hap-mer (haplotype-specific k-mers as defined by Mercury ^71^) databases were generated and the haplotype assemblies evaluated using the scaffolds of both haplotypes. For building of phased blocks, the parameter ‘num_switch 10’ and ‘short_range 20,000’ were applied. For comparison, Merqury was also run on PN40024, on VitRGM, on both Cabernet Sauvignon haplotype assemblies v1.1 and on both *M. rotundifolia* cultivar ‘Trayshed’ haplotype assemblies v2.0 ^23^. The 19-mer profiles were computed from PN40024 WGS reads SRR5627797, from VitRGM WGS reads SRR8379638, from Cabernet Sauvignon WGS reads SRR3346861 and SRR3346862 and from *M. rotundifolia* WGS reads SRR6729333. Prior to analyses, the reads were trimmed with Trimmomatic-v0.39 allowing a minimum read length of 80 or 90 (PN40024 reads).

### Gene annotation

The ‘Börner’ haplotypes were annotated *ab initio* with MAKER v3.01.03 following the MAKER-P pipeline ^72, 73^. As input data, haplotype-specific repeat libraries were created according to the MAKER advanced repeat library protocol. The monocotyledons repeat library of RepBase (RepeatMaskerEdition-20181026, model_org=monocotyledons) ^74^, the transposable element (TE) sequences from MAKER, the ‘Börner’ *de novo* transcriptome assembly described above, *Vitis* protein sequences (NCBI, Protein DB “Vitis”[Organism]), plant protein sequences (UniProt, release 2020_02, “Viridiplantae [33090]” AND reviewed:yes) and *Vitis* full-length cDNAs (NCBI, Nucleotide DB “Vitis” [Organism] AND complete cds[Title]) were used. The gene predictors SNAP v2006-07-28 (three rounds) ^75^ and GeneMark v3.60 ^76^ were trained on both, BoeRip and BoeCin. AUGUSTUS v3.2.3, tRNAscan-SE v1.3.1 and EvidenceModeler v1.1.1 ^77^, SNAP and GeneMark together were ran through MAKER to compute the final gene annotation. The option ‘split_hit’ was set to 20,000 and the previously generated *Vitis* parameter set was adjusted for AUGUSTUS.

The gene models from BoeRip and BoeCin were refined with PASA v2.4.1 ^78^. As evidence data, the *de novo* and reference-guided ‘Börner’ transcriptome assemblies, the *Vitis* full-length cDNAs and *Vitis* EST data (NCBI, txid3603[Organism:exp] AND is_est[filter]) were used. The parameter file is available as Supplementary File 3 Data S1. The refinement with PASA was iteratively applied three times on the gene models. The reference-guided assembly was computed with Trinity v2.10.0 using the ‘Börner’ RNA-Seq data aligned with HISAT2 v2.2.0. Only alignments with MQ >10 were given to Trinity and ‘--genome_guided_max_intron’ was set to 20,000.

After structural gene annotation, a functional gene annotation was added through a BLAST search (package+ v2.10.0) against the UniProt/Swiss-Prot database (release 2020_02), through domain identification with InterProScan5 v5.42-78.0 ^79^ and the PFAM database v32.0. Gene models with Annotation Edit Distance (AED) > 0.5 were filtered from the final gene model set. Also, predicted genes expected to encode a polypeptide with less than 50 amino acids and no functional annotation were discarded. The annotation data for BoeRC are available at (https://doi.org/10.4119/unibi/2962793). For comparison, functional annotation was also carried out for the PN40024 VCost.v3 structural gene annotation with identical resources.

### Prediction of resistance genes

Resistance gene analogs (RGAs) were predicted for both haplotypes with the pipeline RGAugury v2.1.7 ^29^. RGAugury employed ncoil ^80^, PfamScan v1.6 ^81^, InterProScan5 v5.45-80.0 ^79^ and Phobius-1.01 ^82^ on the protein sequences to predict known resistance domains. The initial filtering against the RGAdb was disabled since the database has not been updated since 2016, unknown RGAs not included in the database were of interest and the identification of (resistance) domains for none RGAs was enabled. For InterProScan5 domain search through RGAugury, the databases Gene3D-4.2.0 ^83^, Pfam-33.1 ^84^, SMART-7.1 ^85^ and SUPERFAMILY-1.75 ^86^ were utilized.

The RGA annotations of the two haplotypes were correlated through determination of RBHs based on protein sequences. Only RBHs with an identity and coverage ≥ 90 were considered.

### Assembly wide variant detection

The SNP and InDel positions of the SNP output from the DNAdiff analysis (see above) were compared with the positions of the coding and non-coding gene regions and classified accordingly.

### Genetic mapping

The annotated genes and nucleotide sequences of the phylloxera resistance QTL on chr13, referred to as *Rdv1*, of BoeRip, BoeCin, PN40024 and VitRGM were compared. Before this study, the QTL *Rdv1* was delimited with the associated genetic markers GF13-01 and GF13-12 ^6^; the marker positions were identified based on the respective marker assay primer sequences with primersearch (EMBOSS package ^63^). To narrow down the location of *Rdv1*, additional genetic fine mapping was carried out with a mapping population of in total 498 F1 genotypes (see above) and newly designed sequence-tagged SSR (STS) markers. The additional markers were derived from PN40024 ^16^ (Supplementary File 2 Table S14) and used to screen the F1 genotypes from the mapping population.

Five relevant genotypes with marker data indicating recombination events within the initial *Rdv1* QTL were subjected to GBS to locate the recombination points between the two haplotypes of ‘Börner’. V3125 was included as a control for comparative read mapping. The SE reads were quality trimmed using Trimmomatic-0.39^46^ with the following parameters: ILLUMINACLIP: 4:30:15 LEADING:30 TRAILING:30 SLIDINGWINDOW:4:15 MINLEN:60. Trimmed reads of each of the six studied genotypes as well as reads from ‘Börner’ ^10^ were separately mapped to BoeCin and BoeRip using the HISAT2 version 2.2.0 ^59^ with the following parameters: --no-softclip –no-spliced-alignment. The obtained SAM files were converted to sorted BAM files as well as indexed using SAMtools. Variant calling against BoeRC and estimation of genomic recombination sites was supported by QIAGEN CLC Genomics Workbench 22.0 (QIAGEN, Aarhus, Denmark). SNPs detected in the *Rdv1* region on chr13 for each genotype were filtered with respect to read coverage and frequency. Homozygous variants were counted if > 90 % of the reads support the variation, for heterozygous variants a frequency between 25 % and 75 % was required. An example case (GBS-04) for the assessment of GBS markers is shown in Supplementary File 1 Fig. S15. Recombination sites were determined by monitoring the switch from either the BoeCin haplotype to the BoeRip haplotype or *vice versa* between a set of at least two variant positions.

### Allele sequence comparison and TE detection

The genes (alleles, orthologs) of the *Rdv1* region of BoeRip, BoeCin, PN40024 and VitRGM between the markers GF13-03 and GF13-07 were placed into groups with OrthoFinder v2.3.11 ^87^ using the longest protein sequence annotated for the splicing variants of each gene. Missing alleles (or, alternatively phrased, missing predictions of orthologous genes at syntenic positions) were investigated and re-annotated by aligning the protein sequences of the other cultivars against the genome sequences with exonerate v2.4.0 ^88^. Only alignments with at least 70 % coverage of the protein sequence were considered. To obtain gene annotations for remaining poorly annotated sequences despite existing RNA-Seq support in the genomic region studied, BRAKER was run with strictly filtered RNA-Seq mappings (only uniquely mapped PE reads on both ‘Börner’ haplotypes offered as combined target, both reads of a pair must map); adding untranslated regions (UTRs) and prediction of alternative transcripts from RNA-Seq data was enabled. Since the focus here is on phylloxera resistance at the root, an additional AUGUSTUS species model was trained on BoeRC with the uniquely mapped root RNA-Seq reads, the above described *Vitis* protein and plant protein sequences and eudicot protein sequences from OrthoDB v10.1 ^89^. The results were manually evaluated and restored gene models of BoeRip and BoeCin were refined two times with PASA. Through investigation of the RNA-Seq mappings from all available tissues to BoeCin, the gene predictions in the *Rdv1* region of all four sequence regions studied were manually screened for low supported gene models. Transcripts encoding for protein sequences with less than 50 amino acids, tagged as ‘protein of unknown function’ (PUF) and with no ortholog were removed. RGA classes were determined for the manually curated genes/alleles with RGAugury as described above. TEs were identified with the Extensive *de novo* TE Annotator (EDTA) ^90^. Finally, the results were manually collected in allelic representations of the genes of the *Rdv1* region of BoeRip, BoeCin, PN40024 and VitRGM. An extended version of the gene overview at the *Rdv1* locus that includes TEs has also been prepared (Supplementary File 1 Fig. S11).

## Authors’ contributions

Sampling and phenotyping were done at the JKI (Julius Kuehn Institute, Institute for Grapevine Breeding Geilweilerhof, Siebeldingen, Germany). Sequencing and data analysis were performed at Bielefeld University, Faculty of Biology & Center for Biotechnology (CeBiTec) and MPIPZ (Cologne). The design of the experiments was set up by LH, DH and BF. Genotype selection, sampling and DNA isolations were carried out by LH and DH. PV, BH and RR accomplished library preparation and sequencing. LH performed the SSR analysis. BW supervised the work at Bielefeld University. RT supervised the work at JKI. RT and BW acquired project funding and wrote the project proposal. All bioinformatic data analyses, creation of figures, tables and writing of the initial draft of the manuscript were performed by BF and DH. BF and BW finalised the manuscript. All authors have read and approved the final manuscript.

## Acknowledgements

The authors like to thank all members of the Genetics and Genomics of Plants team at Bielefeld University as well as the members of the Julius Kuehn Institute for Grapevine Breeding at Geilweilerhof.

The project was supported by funds from the Federal Ministry of Food and Agriculture (BMEL), based on a decision by the Parliament of the Federal Republic of Germany via the Federal Office for Agriculture and Food (BLE) under the innovation support program (project acronym MureViU, 28-1-82.066-15), as well as by the EU COST Action INTEGRAPE (CA 17111). We acknowledge support for the article processing charge by the Deutsche Forschungsgemeinschaft and the Open Access Publication Fund of Bielefeld University.

This work was supported by the BMBF-funded de.NBI Cloud within the German Network for Bioinformatics Infrastructure (de.NBI).

## Data availability

The SMRT sequencing reads of ‘Börner’ (ERR6182799), the WGS reads of ‘Börners’ parents *V. riparia* GM18 (ERR6182967, ERR6183009) and *V. cinerea* Arnold (ERR6183041) as well as the ‘Börner’ haplotype assemblies (ERS6637871) were deposited in ENA/GenBank/DDJ under project no. PRJEB45595. The haplotype assemblies BoeRip and BoeCin at chromosome level with structural and functional annotation are available at (https://doi.org/10.4119/unibi/2962793). The RNA-Seq data of ‘Börner’ leaf (ERR6182801) and tendril (ERR6182801) were deposited in ENA/GenBank/DDJ under project no. PRJEB46079, the raw sequencing reads of ‘Börner’ BACs (ERR6183016-ERR6183021, ERR6183023-ERR6183025, ERR6183030-ERR6183032) under project no. PRJEB46081. The assembled and phase-separated BAC contigs are available at (https://doi.org/10.4119/unibi/2962639). DNA sequence read data from the F1 genotypes of the mapping population and the maternal parent V3125 generated for GBS were deposited in ENA/GenBank/DDJ under project no. PRJEB53997.

## Supplementary information

Supplementary File 1 Fig. S1: Cumulative length of both ‘Börner’ haplotype assemblies

Supplementary File 1 Fig. S2: Sequence region with BAC sequences of both haplo-types

Supplementary File 1 Fig. S3: Spectra-copy number and assembly spectrum plot of the ‘Börner’ genome sequence assembly

Supplementary File 1 Fig. S4: Analysis of phasing with Mercury

Supplementary File 1 Fig. S5: All-versus-all dot plot between pseudochromosomes of the *V. riparia* haplotype of ‘Börner’ and the *V. vinifera* reference sequence.

Supplementary File 1 Fig. S6: All-versus-all dot plot between pseudochromosomes of the *V. cinerea* haplotype of ‘Börner’ and the *V. vinifera* reference sequence

Supplementary File 1 Fig. S7: All-versus-all dot plot between pseudochromosomes of the *V. riparia* haplotype of ‘Börner’ and those of *V. riparia* Gloire de Montpellier

Supplementary File 1 Fig. S8: All-versus-all dot plot between pseudochromosomes of the *V. cinerea* haplotype of ‘Börner’ and those of *V. riparia* Gloire de Montpellier

Supplementary File 1 Fig. S9: Dot plot of the extended *Rdv1* sequence region on pseudochromosome 13 between BoeRip and PN40024

Supplementary File 1 Fig. S10: Dot plot of *Rdv1* on pseudochromosome 13 between BoeCin and PN40024

Supplementary File 1 Fig. S11: Extended version of Fig. 6, “Genes of BoeCin, BoeRip, PN40024 and VitRGM at and surrounding the *Rdv1* locus”, transposable elements/TEs included

Supplementary File 1 Fig. S12: Length distribution calculated over all subreads

Supplementary File 1 Fig. S13: Coverage plot of the ‘Börner’ haplotype assemblies

Supplementary File 1 Fig. S14: Display of initial assembly analysis of chr02 of BoeRip

Supplementary File 1 Fig. S15: Example for determination of allelic constitution in selected F1 genotypes

Supplementary File 2 Table S1: Pseudochromosome sizes of the haplotype assemblies BoeRip and BoeCin

Supplementary File 2 Table S2: Quality statistics of *Vitis* genome assemblies

Supplementary File 2 Table S3: Assembly statistics of the BAC contigs

Supplementary File 2 Table S4: Mapping statistics of the BAC contigs on the BoeRip and BoeCin haplotype assembly

Supplementary File 2 Table S5: Statistics of the trimmed BAC sequences

Supplementary File 2 Table S6: Haplotype blocks of BoeRip and BoeCin

Supplementary File 2 Table S7: Repeat content of the haplotypes

Supplementary File 2 Table S8: GBS marker sites for the parents ‘Börner’ and V3125 and selected F1 offspring

Supplementary File 2 Table S9: BoeCin, BoeRip, PN40024 and VitRGM genes in *Rdv1* region

Supplementary File 2 Table S10: Amino acid sequence similarity matrix BoeCin to BoeCin deduced from genes in *Rdv1* region

Supplementary File 2 Table S11: Amino acid sequence similarity matrix BoeCin to BoeRip deduced from genes in *Rdv1* region

Supplementary File 2 Table S12: Amino acid sequence similarity matrix BoeCin to PN40024 deduced from genes in *Rdv1* region

Supplementary File 2 Table S13: Amino acid sequence similarity matrix BoeCin to VitRGM deduced from genes in *Rdv1* region

Supplementary File 2 Table S14: Primer sequences for SSR marker

Supplementary File 2 Table S15: RNA-Seq read trimming statistics

Supplementary File 2 Table S16: Trimming statistics of *V. riparia* and *V. cinerea* genomic reads

Supplementary File 2 Table S17: Description of samples applied to GBS

Supplementary File 2 Table S18: Mapping positions of the genetic SSR marker on the PN40024 genome assembly and on the ‘Börner’ haplotype assemblies BoeRip and BoeCin

Supplementary File 2 Table S19: Distribution of the 344 genetic marker used for pseudochromosome validation over BoeRip and BoeCin.

Supplementary File 3 Data S1: Parameter file for PASA gene model refinement

