## Supplementary File 1 for "A fully phased interspecific grapevine rootstock genome sequence representing *V. riparia* and *V. cinerea* and allele-aware annotation of the phylloxera resistance locus *Rdv1*"

### Supplementary Figures

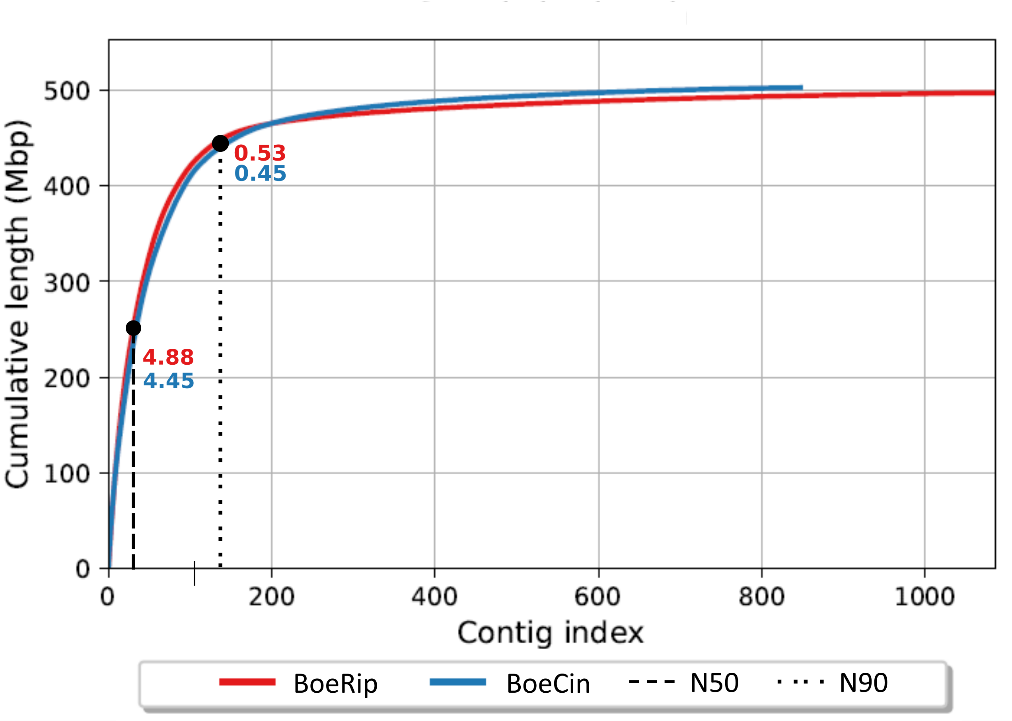

#### Fig. S1 – Cumulative length of both ‘Börner’ haplotype assemblies.

The plot displays the cumulative length of the contigs of the ‘Börner’ haplotype assemblies. The red line represents the BoeRip haplotype assembly and the blue line the BoeCin haplotype assembly. The N50 and N90 values are marked and given in [Mbp].

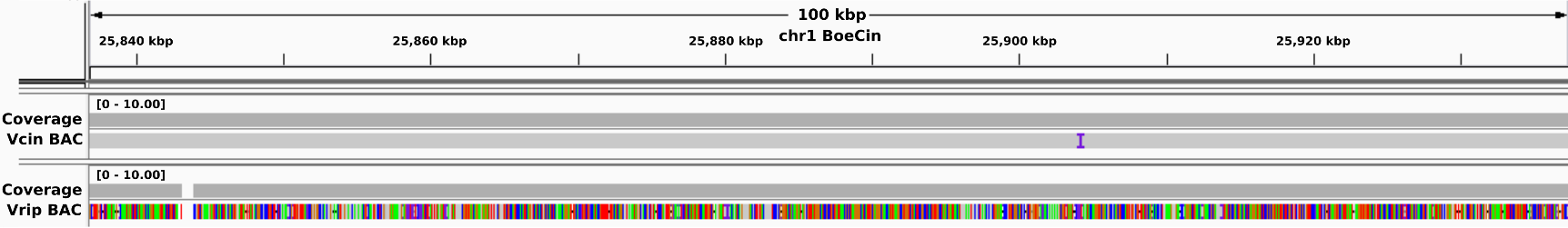

#### Fig. S2 – Sequence region with BAC sequences of both haplotypes.

The figure displays a sequence region on chr01 of the BoeCin haplotype on which 'Börner' BAC sequences from both phases map. Here, the BAC sequence from the V. cinerea haplotype of 'Börner' (Vcin BAC, gray bar with one purple mark) maps with only one insertion, while the BAC sequence from the V. riparia haplotype of 'Börner' (Vrip BAC, fully coloured bar) maps with plenty of mismatches and indels.

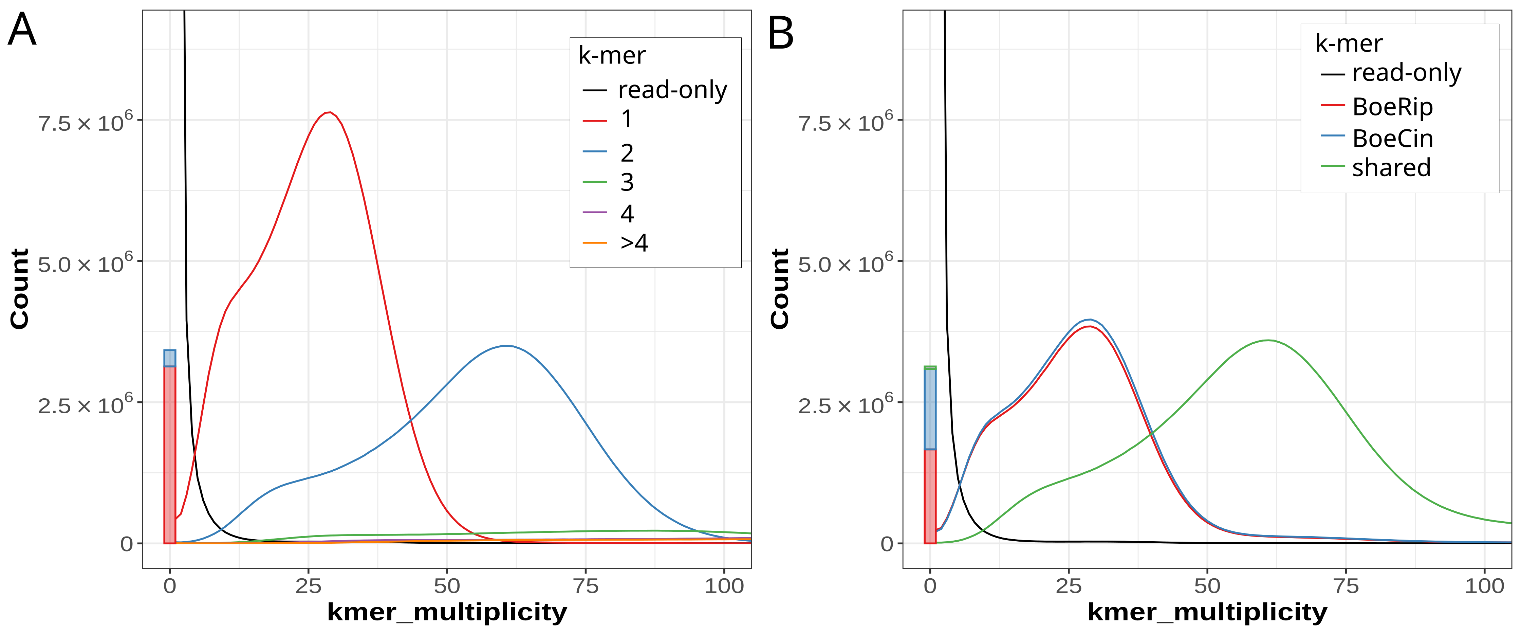

#### Fig. S3 – Spectra-copy number and assembly spectrum plot of the ‘Börner’ genome sequence assembly.

Black read-only area refers to k-mers found only in the ‘Börner’ WGS reads while they are not present in the diploid sequence assembly BoeRC. Almost no read-only (missing) k-mers were detected. (A) The left panel displays the spectra-copy number plot. A higher 1-copy peak (red peak) compared to the 2-copy peak (blue peak) was detected. The 1-copy peak likely refers to k-mers derived from single-copy regions (expected heterozygous regions between BoeRip and BoeCin) and the 2-copy peak to k-mers derived from sequences present in both haplotypes (expected homozygous regions). (B) The right panel displays the assembly spectrum plot showing the shared amount of k-mers (area below green line) present in both haplotypes. The mostly overlapping red and blue lines indicate the k-mers unique to either BoeRip or BoeCin.

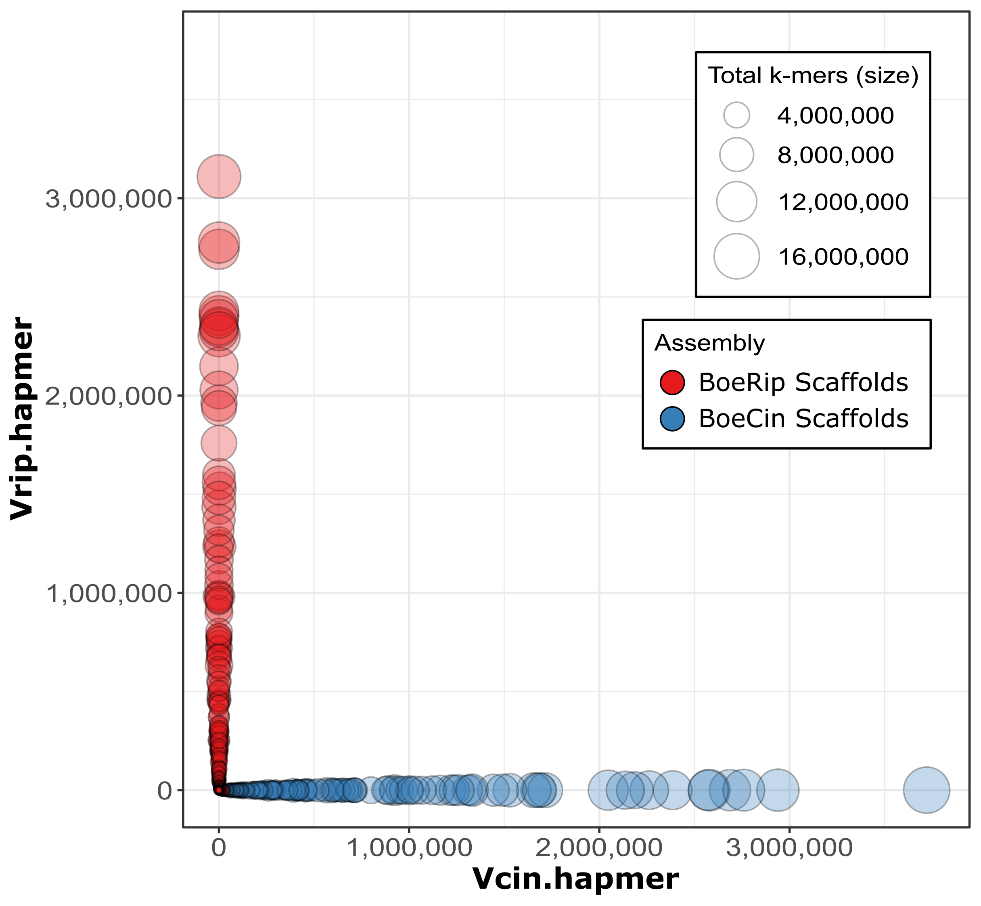

#### Fig. S4 – Analysis of phasing with Mercury.

Each circle represents a scaffold sequence of the BoeRip (red) or BoeCin (blue) haplotype assembly. The x-axis shows the number of V. cinerea specific 19-mers (Vcin.hapmer) and the y-axis the number of V. riparia specific 19-mers (Vrip.hapmer).

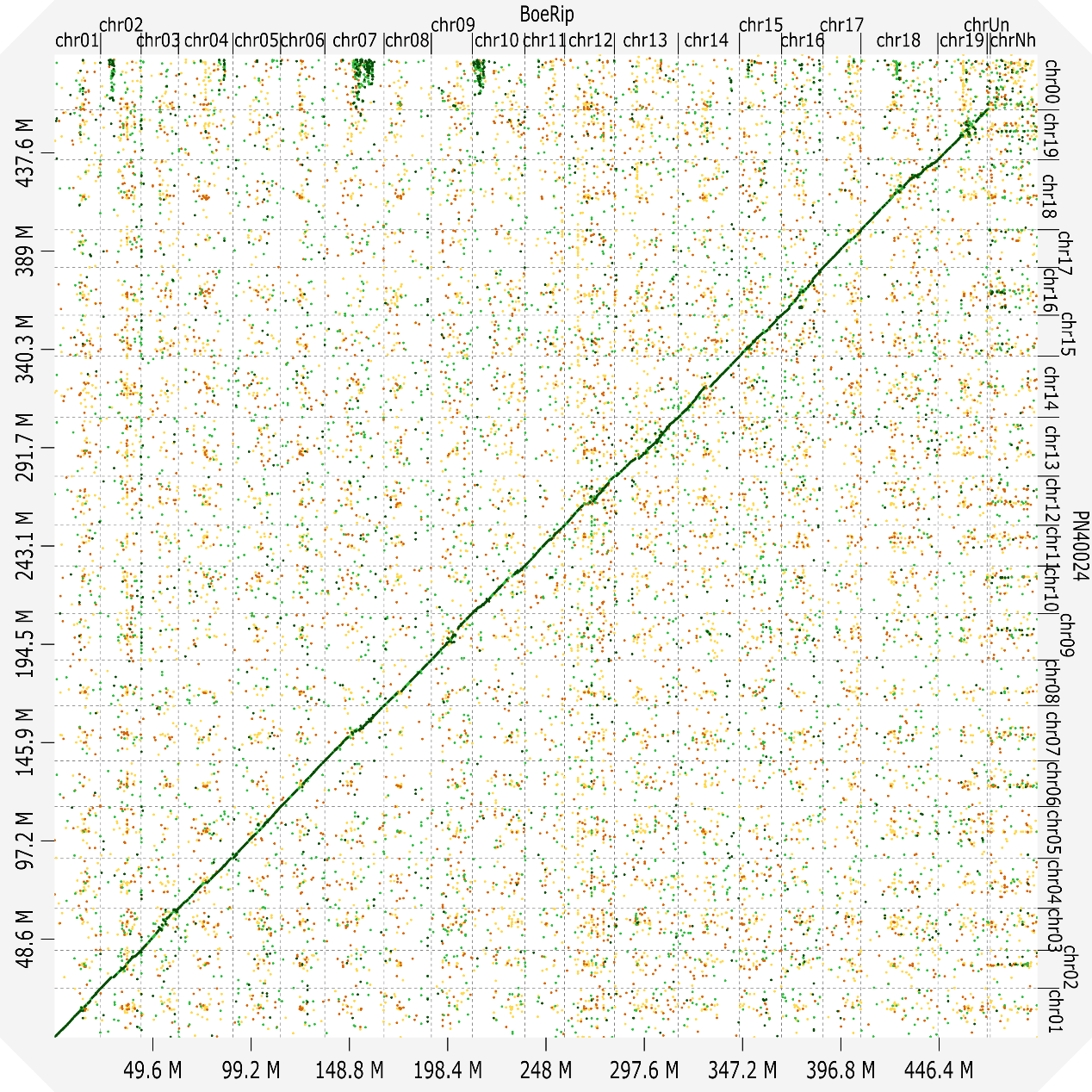

#### Fig. S5 – All-versus-all dot plot between pseudochromosomes of the *V. riparia* haplotype of 'Börner' and the *V. vinifera* reference sequence.

The graphic shows the dot plot between the pseudochromosomes of BoeRip and PN40024.

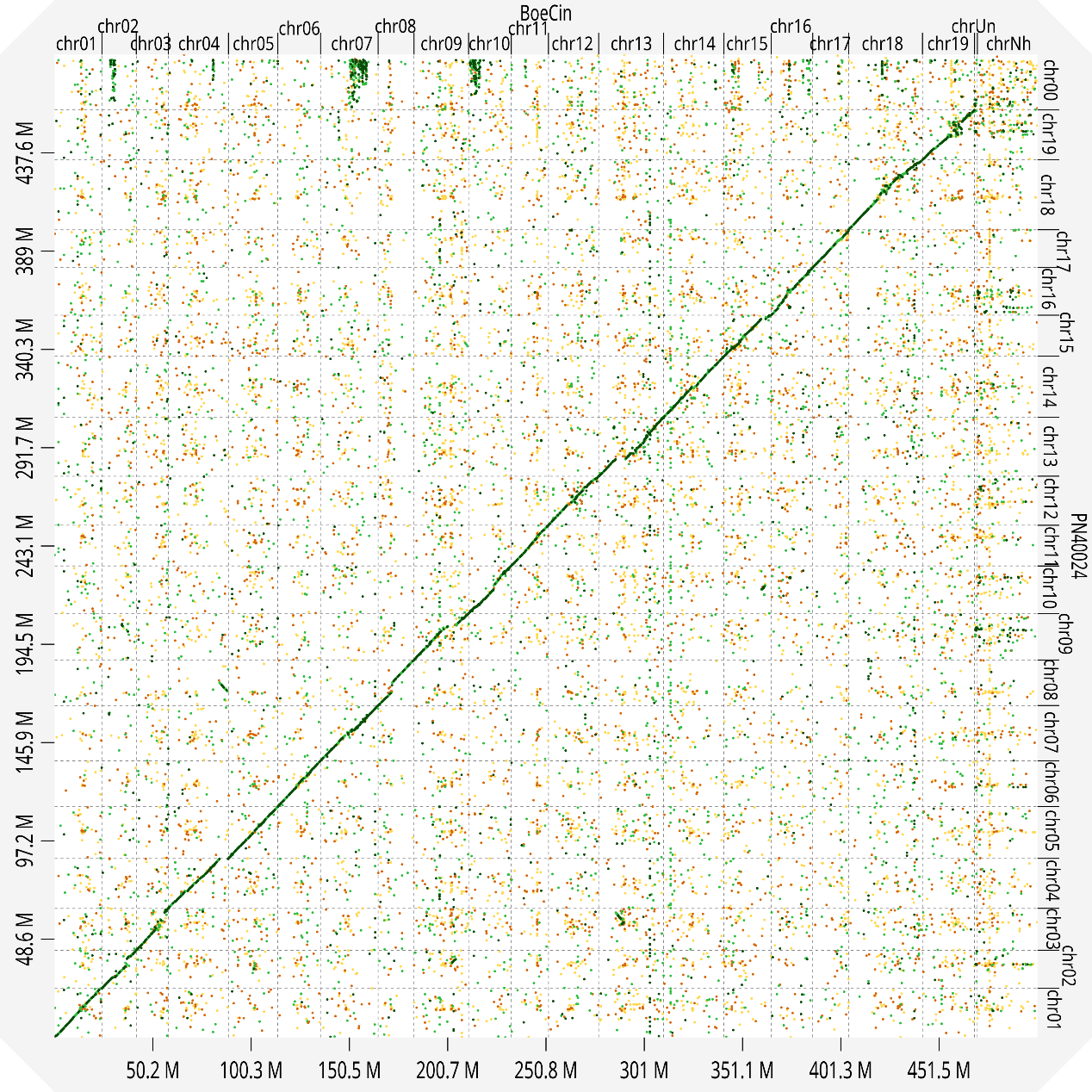

#### Fig. S6 – All-versus-all dot plot between pseudochromosomes of the V. *cinerea* haplotype of 'Börner' and the *V. vinifera* reference sequence.

The graphic shows the dot plot between the pseudochromosomes of BoeCin and PN40024. The hint for the potential mis-assembly mentioned in the main text is visible in the lower left region (chr04 from left and chr08 from bottom).

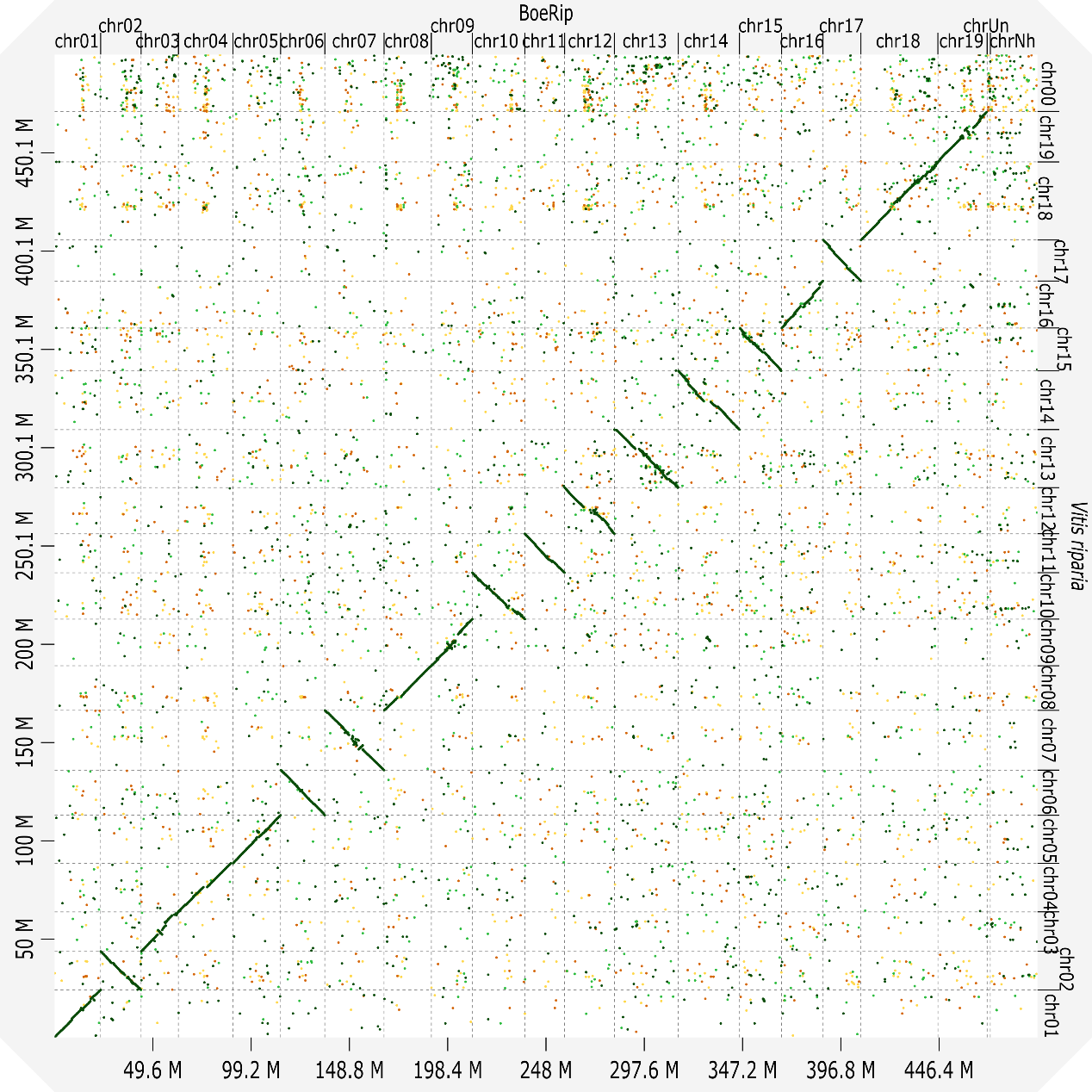

#### Fig. S7– All-versus-all dot plot between pseudochromosomes of the V. *riparia* haplotype of 'Börner' and those of *V. riparia* Gloire de Montpellier.

The graphic shows the dot plot between the pseudochromosomes of BoeRip and VitRGM. Note that in VitRGM some pseudochromosomes are included in reverse orientation.

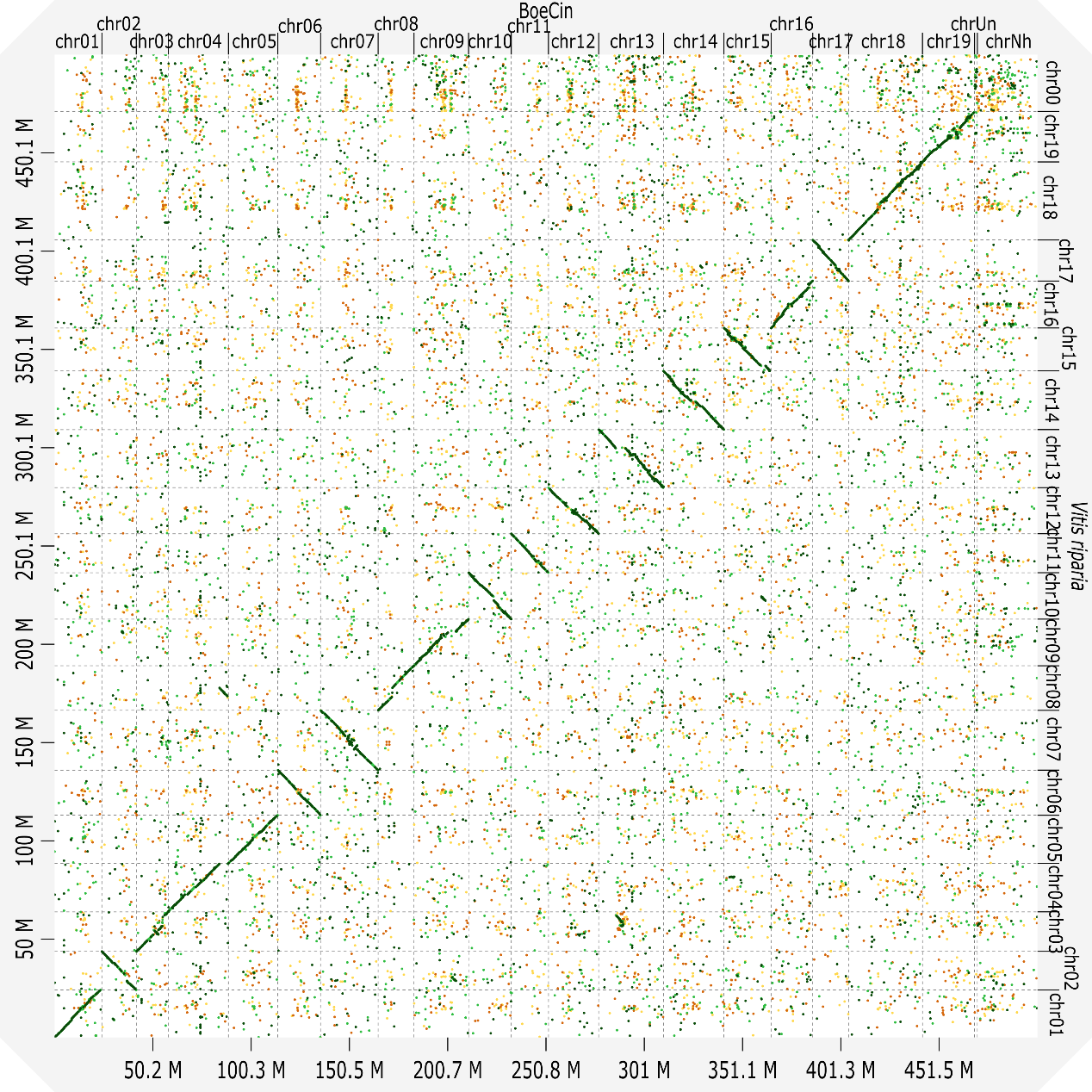

#### Fig. S8 – All-versus-all dot plot between pseudochromosomes of the V. *cinerea* haplotype of 'Börner' and those of *V. riparia* Gloire de Montpellier.

The graphic shows the dot plot between the pseudochromosomes of BoeCin and VitRGM. Note that in VitRGM some pseudochromosomes are included in reverse orientation.

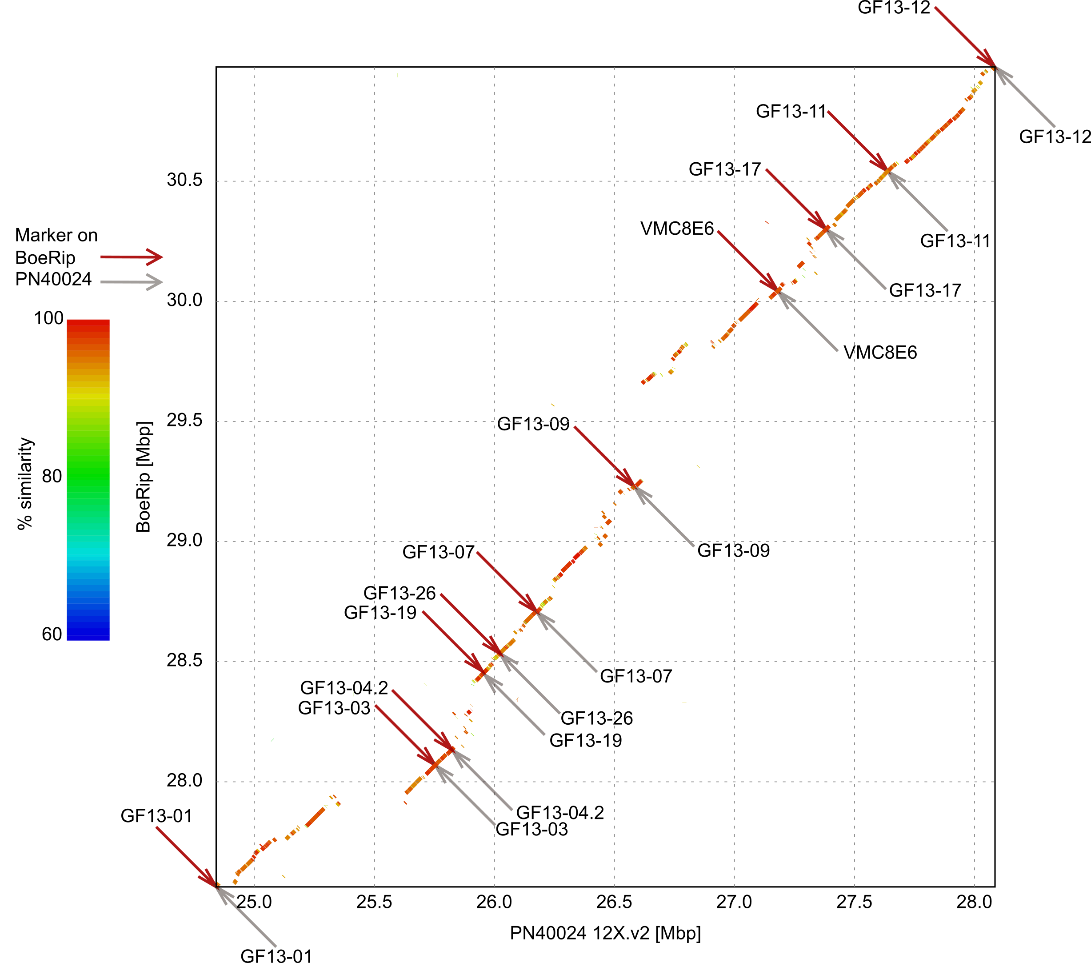

#### Fig. S9 – Dot plot of the extended *Rdv1* sequence region on pseudochromosome 13 between BoeRip and PN40024.

Genetic markers mapping to BoeRip are represented as red arrows, genetic markers mapping to PN40024 as grey arrows.

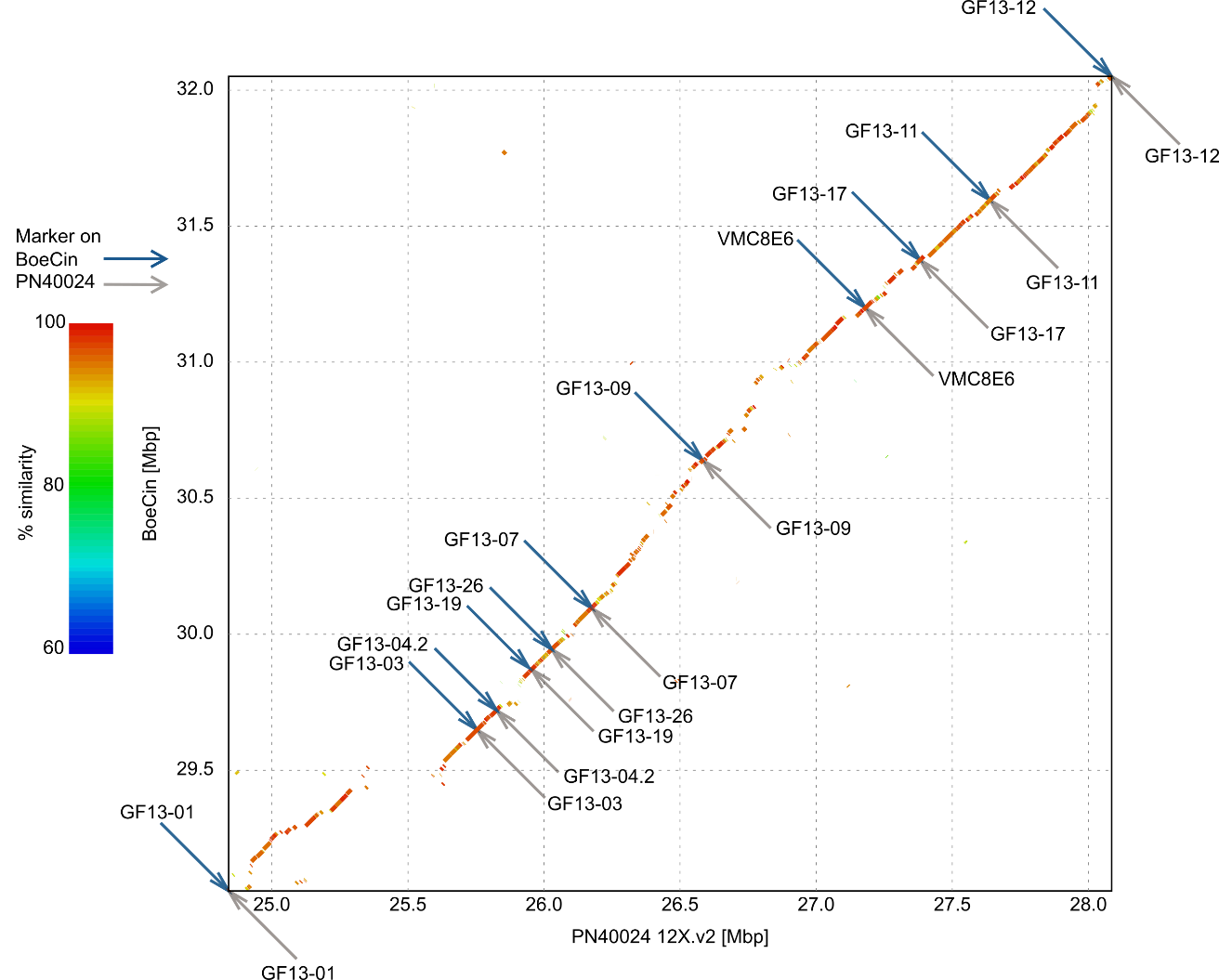

#### Fig. S10 – Dot plot of *Rdv1* on pseudochromosome 13 between BoeCin and PN40024.

Genetic markers mapping to BoeCin are represented as blue arrows, genetic markers mapping to PN40024 as grey arrows.

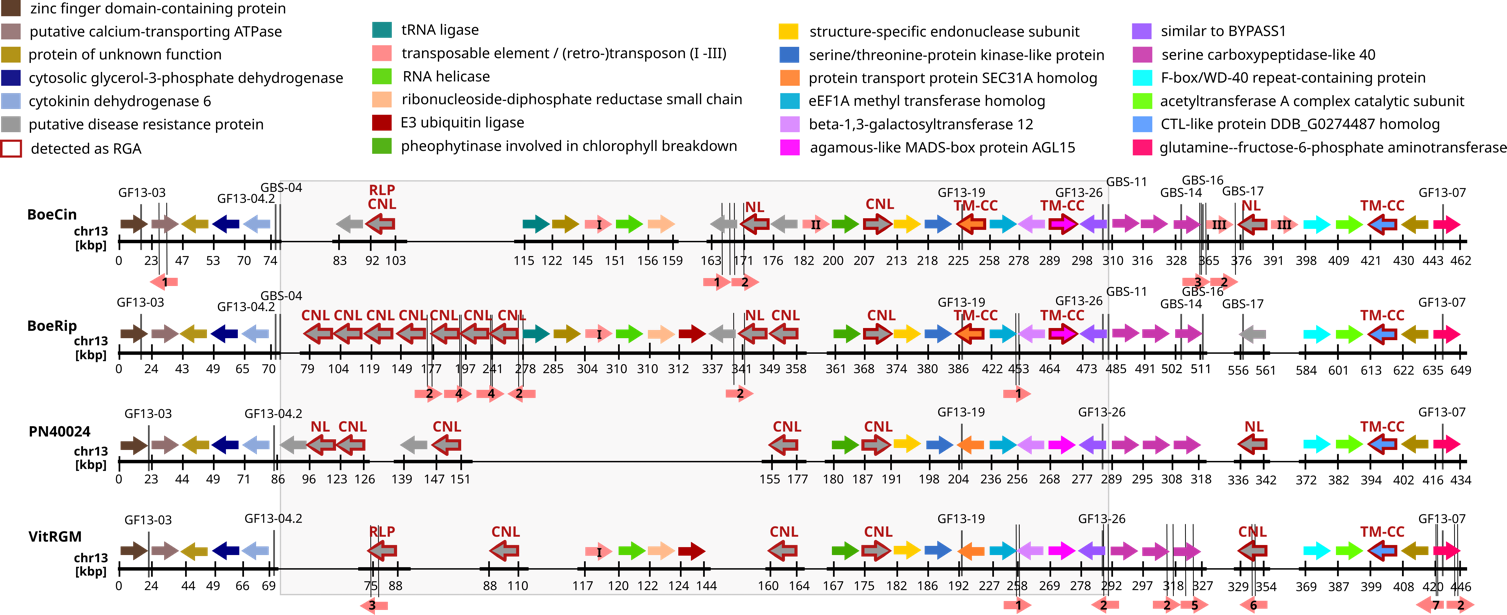

#### Fig. S11 – Extended version of Fig. 6, "Genes of BoeCin, BoeRip, PN40024 and VitRGM at and surrounding the *Rdv1* locus", transposable elements/TEs without annotated gene structures included.

The genes (alleles) were clustered according to similarity and are shown in one colour for each orthogroup. The grey box represents the Rdv1 locus delineated by the genetic markers GF13-04.2/GBS-04 and GF13-26/GBS-11; additional markers are included upstream and downstream of the Rdv1 locus. The kbp values on the axis were adapted such that the start of the first gene is bp zero (relative coordinates). At the top, the colour code for each group of related genes/alleles and the corresponding functional annotation is given. Grey coloured genes are potential resistance genes according to their functional annotation; genes with a red edging were identified as RGA (see text). Above each RGA, the domain type classification of the encoded protein is mentioned. See legend to Fig. 4 for acronyms, CC-NBS-LRR abbreviated as CNL, NBS-LRR as NL. Three genes qualified as TE genes of different types, these are marked with roman numbers; I: mutator-like element (MULE), II: similar to transposon TX1 protein, III: similar to retrovirus-related polymerase polyprotein.
In the lower part of each track, TE positions are indicated by rose coloured arrows. Only intact or almost complete transposons are shown, TE fragments are excluded. Class I (retrotransposons) and class II (DNA transposons) TEs are not distinguished. The arrows with arabic numbers are not drawn to scale; the TEs might be located within genes or in intergenic regions; 1: LTR/Gypsy, 2: LTR/Copia, 3: DNA/DTC, 4: DNA/DTA (hAT), 5: DNA/Helitron, 6: MITE/DTM, 7: DNA/DTM. Designations taken from Extensive de novo TE Annotator (EDTA).

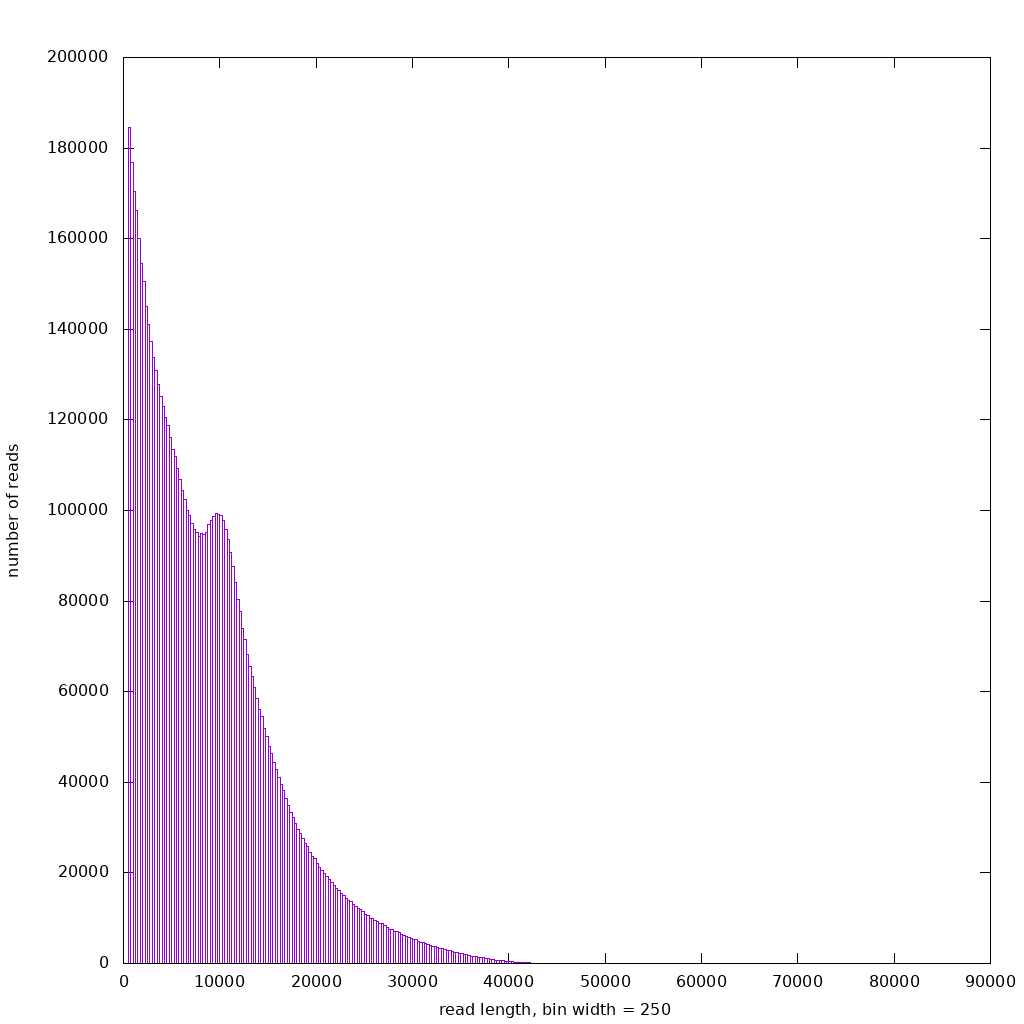

.

#### Fig. S12 - Length distribution calculated over all subreads.

The length distribution of all SMRT subreads of the raw sequencing data-set. The graph was computed with Canu.

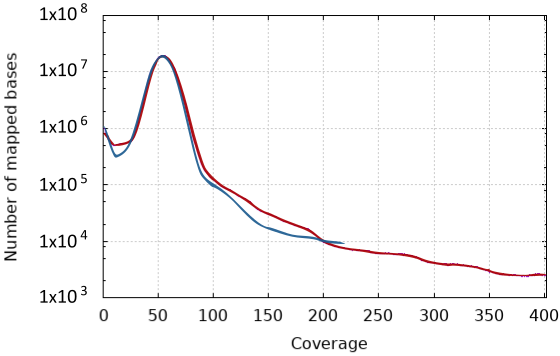

#### Fig. S13 – Coverage plot of the ‘Börner’ haplotype assemblies.

The plot shows the coverage per number of mapped bases of the haplotype assemblies. The red line represents the BoeRip haplotype assembly and the blue line the BoeCin haplotype assembly.

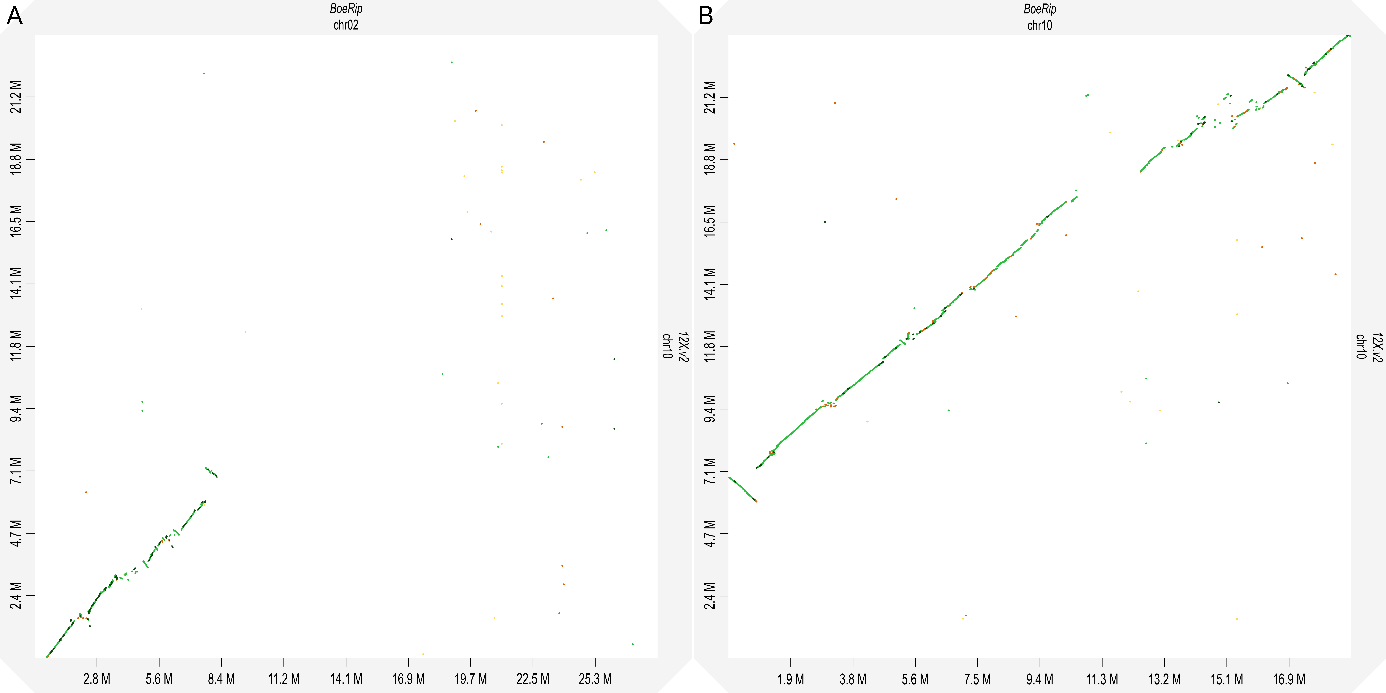

#### Fig. S14 – Display of initial assembly analysis of chr02 of BoeRip.

**(A)** The dot plot in the left panel shows similarity between pseudochromosome 02 of BoeRip and pseudochromosome 10 of PN40024 that was detected initially and before correction; **(B)** shows a dot plot between pseudochromosome 10 of BoeRip and pseudochromosome 10 of PN40024. The sequence region detected in (A) fits into the gap of (B).

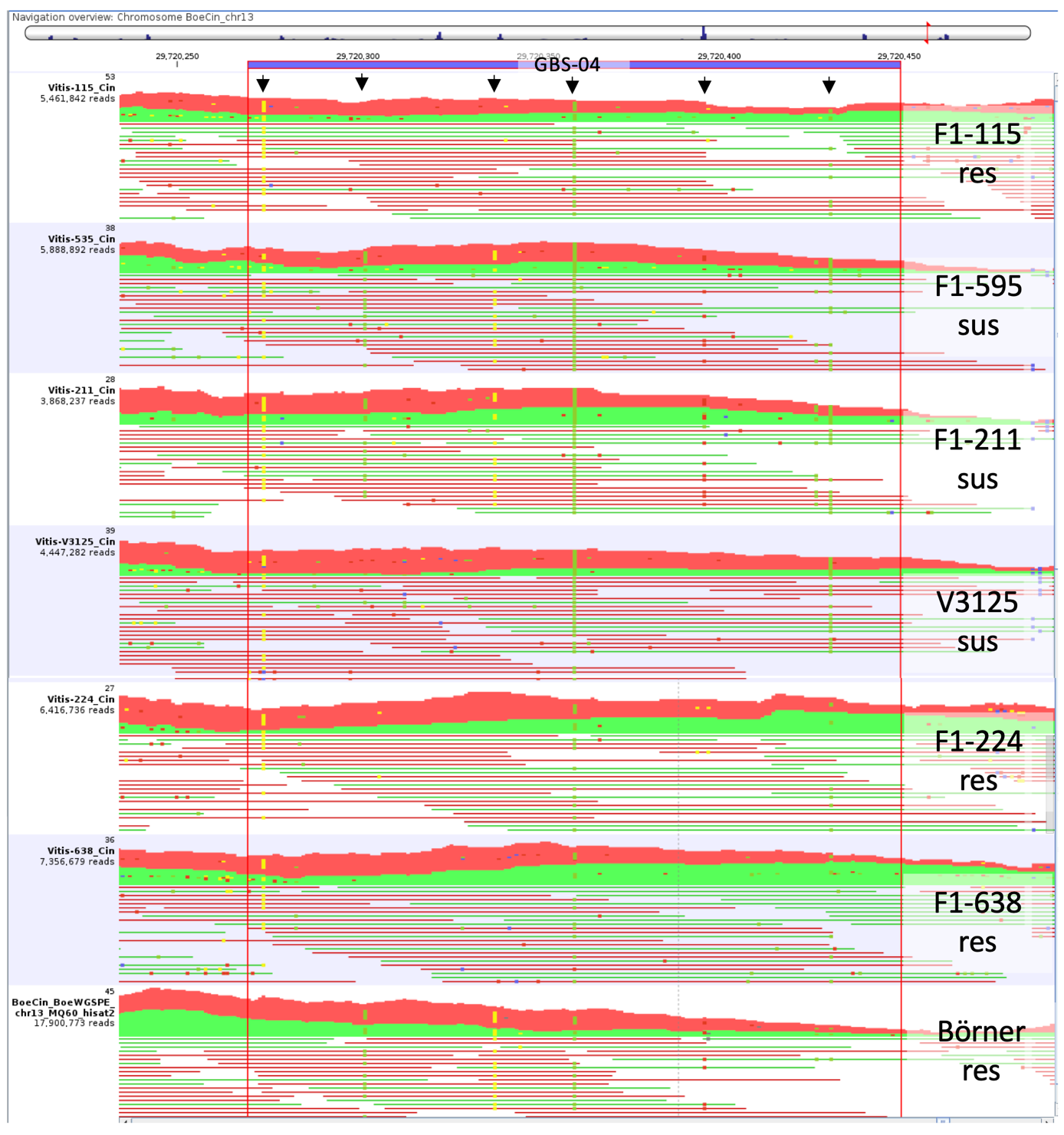

#### Fig. S15 – Example for determination of allelic constitution in selected F1 genotypes.

Shown are mapped reads and SNVs of parental lines and selected F1 individuals on BoeCin (chr13 around the GBS-04 marker). Six SNVs are marked with an arrow, the green and red lines represent individual forward and reverse reads, respectively. Genotype designations and their resistant and susceptible phenotypes are shown on the right.
The GBS-04 marker is heterozygous in 'Börner' (C/T for BoeCin/BoeRip) and homozygous in V3125 (T/T), therefore the susceptible genotypes are homozygous like V3125 even if they contain the BoeRip allele (T/T), whereas the resistant lines inherited the BoeCin-allele and are heterozygous at this position (C/T).
Mismatch A is shown in red, mismatch C is blue, mismatch G is yellow and mismatch T is green.
Numbers on the left of each track are the average number of reads mapped per position in the respective window and the total number of reads mapped to pseudochromosome 13.
