## Supplementary File 3 for "A fully phased interspecific grapevine rootstock genome sequence representing *V. riparia* and *V. cinerea* and allele-aware annotation of the phylloxera resistance locus *Rdv1*"

### Supplementary Data

**Data S1** **– Parameter file for PASA gene model refinement:**

validate_alignments_in_db.dbi:--MIN_PERCENT_ALIGNED=90

validate_alignments_in_db.dbi:--MIN_AVG_PER_ID=80

validate_alignments_in_db.dbi:--MAX_INTRON_LENGTH=20000

validate_alignments_in_db.dbi:--NUM_BP_PERFECT_SPLICE_BOUNDARY=0

subcluster_builder.dbi:-m=50

cDNA_annotation_comparer.dbi:--MIN_PERCENT_OVERLAP=50

cDNA_annotation_comparer.dbi:--MIN_PERCENT_PROT_CODING=30

cDNA_annotation_comparer.dbi:--MIN_PERID_PROT_COMPARE=50

cDNA_annotation_comparer.dbi:--MIN_PERCENT_LENGTH_FL_COMPARE=70

cDNA_annotation_comparer.dbi:--MIN_PERCENT_LENGTH_NONFL_COMPARE=70

cDNA_annotation_comparer.dbi:--MIN_PERCENT_ALIGN_LENGTH=70

cDNA_annotation_comparer.dbi:--MIN_PERCENT_OVERLAP_GENE_REPLACE=90

cDNA_annotation_comparer.dbi:--MAX_UTR_EXONS=5
